# N*-*acetyl-transferases required for iron uptake and aminoglycoside resistance promote virulence lipid production in *M. marinum*

**DOI:** 10.1101/2024.07.05.602253

**Authors:** Bradley S. Jones, Vikram Pareek, Daniel D. Hu, Simon D. Weaver, Camille Syska, Grace Galfano, Matthew M. Champion, Patricia A. Champion

**Affiliations:** Department of Biological Sciences, University of Notre Dame, Notre Dame IN, 46556; Eck Institute for Global Health, University of Notre Dame, Notre Dame IN, 46556; Department of Chemistry and Biochemistry, University of Notre Dame, Notre Dame IN, 46556; CNRS – CRBM, Montpellier, France

**Keywords:** Mycobacterium, PDIM, GNAT, Acetylation, MbtK, Eis

## Abstract

Phagosomal lysis is a key aspect of mycobacterial infection of host macrophages. Acetylation is a protein modification mediated enzymatically by N-acetyltransferases (NATs) that impacts bacterial pathogenesis and physiology. To identify NATs required for lytic activity, we leveraged *Mycobacterium marinum,* a nontubercular pathogen and an established model for *M. tuberculosis. M. marinum* hemolysis is a proxy for phagolytic activity. We generated *M. marinum* strains with deletions in conserved NAT genes and screened for hemolytic activity. Several conserved lysine acetyltransferases (KATs) contributed to hemolysis. Hemolysis is mediated by the ESX-1 secretion system and by phthiocerol dimycocerosate (PDIM), a virulence lipid. For several strains, the hemolytic activity was restored by the addition of second copy of the ESX-1 locus. Using thin-layer chromatography (TLC), we found a single NAT required for PDIM and phenolic glycolipid (PGL) production. MbtK is a conserved KAT required for mycobactin siderophore synthesis and virulence. Mycobactin J exogenously complemented PDIM/PGL production in the Δ*mbtK* strain. The Δ*mbtK M. marinum* strain was attenuated in macrophage and *Galleria mellonella* infection models. Constitutive expression of either *eis* or *papA5,* which encode a KAT required for aminoglycoside resistance and a PDIM/PGL biosynthetic enzyme, rescued PDIM/PGL production and virulence of the Δ*mbtK* strain. Eis N-terminally acetylated PapA5 *in vitro*, supporting a mechanism for restored lipid production. Overall, our study establishes connections between the MbtK and Eis NATs, and between iron uptake and PDIM and PGL synthesis in *M. marinum*. Our findings underscore the multifunctional nature of mycobacterial NATs and their connection to key virulence pathways.

**Significance Statement:** Acetylation is a modification of protein N-termini, lysine residues, antibiotics and lipids. Many of the enzymes that promote acetylation belong to the GNAT family of proteins. *M. marinum* is a well-established as a model to understand how *M. tuberculosis* causes tuberculosis. In this study we sought to identify conserved GNAT proteins required for early stages of mycobacterial infection. Using *M. marinum,* we determined that several GNAT proteins are required for the lytic activity of *M. marinum.* We uncovered previously unknown connections between acetyl-transferases required for iron uptake and antimicrobial resistance, and the production of the unique mycobacterial lipids, PDIM and PGLOur data support that acetyl-transferases from the GNAT family are interconnected, and have activities beyond those previously reported.

## Introduction

Acetylation is the movement of an acetyl group from an acetyl donor to an amine including N^α^-amino groups (N-terminal acetylation), the ε-amine of lysine (N^ε^-acetylation) or primary amines of nucleotides, peptides, and small molecules (1). Acetylation can be facilitated enzymatically by N-acetyltransferases (NATs). N-terminal protein acetylation is considered irreversible because N-terminal deacetylases have not been identified. Lysine acetylation is reversible by lysine deacetylases (2–4). Protein acetylation regulates protein folding, stability, localization, and protein-protein interactions (5–11).

*Mycobacterium tuberculosis* is the cause of the human disease tuberculosis (12). In 2022, *M. tuberculosis* infected 10.6 million individuals, resulting in an estimated 1.3 million deaths worldwide (13). *M. marinum* causes a tuberculosis-like disease in poikilothermic fish, and is an occasional human pathogen.

*M. marinum* serves as a model for several aspects of *M. tuberculosis* pathogenesis (14, 15). We previously reported the N-terminal acetylomes of *M. tuberculosis* and *M. marinum* (*16*). We found that 15% of *M. tuberculosis* proteins and 11% of *M. marinum* proteins are acetylated at their N-termini (16). Approximately 16% of the mycobacterial proteome is acetylated at lysine residues (17). Acetylation mapped to metabolism, antibiotic resistance, iron uptake, and lipid synthesis pathways critical to pathogenesis (17–20).

The majority of predicted and established NATs belong to the GCN-5 Related N-acetyltransferase (GNAT) superfamily of enzymes (1). *M. tuberculosis* and *M. marinum* share 23 highly conserved acetyltransferases (16), several of which are capable of acet-ylating more than one type of target (21–31). Substrates of NATs cannot be predicted by protein sequence alone, requiring that the substrates be determined empirically (*3*).

Early following infection, mycobacterial pathogens are taken up by professional phagocytes (32, 33). Following adaptation to the phagosome (34, 35), pathogenic mycobacteria use the ESAT-6-system-1 (ESX-1) Type VII secretion system to lyse the phagosomal membrane and promote interaction with the macrophage cytoplasm (36–40). Several mycobacterial lipids, including phthiocerol dimycocerosate (PDIM) and phenolic glycolipid (PGL), promote optimal phagosomal lysis (41–43), likely because these lipids are required for the wild-type levels of ESX-1 effector proteins (44). Several ESX-1 proteins (EsxA, EspA, EspB, EspC, EspL, EccA_1_, EccB_1_, EccCb_1_, EccE_1_, EsxH, and EspB), and PDIM/PGL biosynthesis proteins (Mas, PpsA, PpsE, Pks15/1, PapA5 and DrrA) are N-terminally or lysine acetylated (16, 17, 45). We recently identified the Emp1 (MMAR_1839) NAT as the sole enzyme that promotes N-terminal acetylation of EsxA in *M. marinum in vivo* (*46*). Here, we sought to identify highly conserved GNAT proteins between *M. marinum* and *M. tuberculosis* that promote virulence through the ESX-1 and PDIM pathways, leveraging genetics, biochemistry, proteomics, functional assays and models of infection.

## Results

### Several GNAT proteins contribute to *M. marinum* hemolytic activity

We generated *M. marinum* strains lacking individual genes with predicted or established NAT activity [Table 1, (46)], and the corresponding complementation strains. We confirmed each strain using PCR (Figure S1) followed by targeted DNA sequencing. ESX-1 and PDIM/PGL are required for optimal phagolytic activity of *Mycobacterium* (42, 47, 48). We tested the requirement of 19 conserved NATs for *M. marinum* hemolytic activity, which can serve as a proxy for phagolytic activity. Six NATs significantly impacted *M. marinum* hemolytic activity (Fig. 1A), while the remaining 13 NATs did not significantly impact hemolysis (Fig. S2A). WT *M. marinum* lysed red blood cells (RBCs), while the Δ*eccCb_1_*strain, which lacks a functional ESX-1 secretion system (37, 49), failed to lyse RBCs (*P*<0.0001 relative to the WT strain) similar to the cell-free control (PBS). None of the *M. marinum* strains lacking individual NATs phenocopied the Δ*eccCb_1_* strain (Figs. 1A and S2A). However, the deletion of several individual NAT genes significantly reduced hemolysis compared to the WT strain, including *argA, rimI*, *pat, MMAR_3205, MMAR_0332* and *mbtK* (all *P<*0.0001 vs the WT strain, Fig. 1A). The Δ*eis* strain was significantly more hemolytic than the WT strain (*P=*0.0032). For all strains, except Δ*pat,* expression of the NAT gene of interest from the mycobacterial optimal promoter significantly increased hemolysis compared to the deletion strain. From these data, we conclude that several putative and established NATs are required for wild-type hemolytic activity in *M. marinum*.

**Figure 1.**
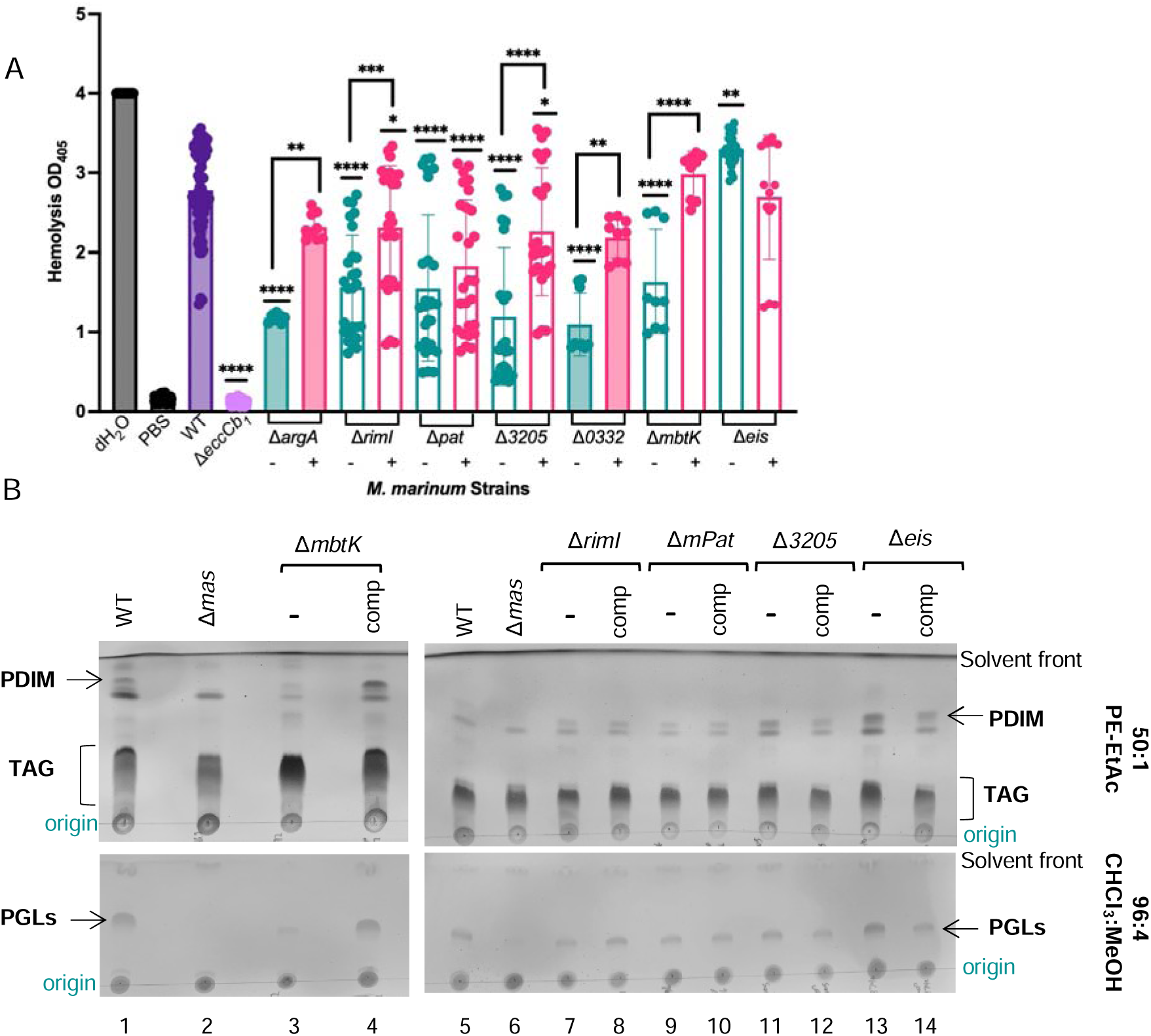
At least four Lysine acetyltransferases are required for WT levels of hemolytic activity in *M. marinum.* A. Hemolytic activity of *M. marinum* strains lacking genes encoding predicted NATs. The putative/ known KATs are not filled. -, + refers to absence or presence of complementation plasmid, which gene of interest expressed from the mycobacterial optimal promoter. Significance relative to the WT strain; Brackets report significance between the deletion and complemented strains. **** *P*<0.0001, *** *P*=0.0004, 0.0002, * *P*=0.0250, 0.0298, ** *P=*0.0032 **B.** TLC analysis of total lipids from the indicated *M. marinum* strains. 6 µl of total lipids were analyzed in each experiment. The solvents used to visualize the indicated lipid species are indicated. The results are representative of at least three biological replicates.

**Table 1.** Table of the *M. marinum* genes with their orthologs from *M. tuberculosis* and their commonly used names.

| *M. marinum* gene | *M. tuberculosis* gene | Name |
| --- | --- | --- |
| MMAR_1067 | Rv0730 |  |
| MMAR_1968 | Rv2747 | ArgA |
| MMAR_4519 | Rv0995 | RimJ |
| MMAR_1839 | Rv2867c | Emp1 |
| MMAR_1882 | Rv2851c | ElaA |
| MMAR_2463 | Rv1653 | ArgJ |
| MMAR_0522 | Rv0262c | Aac |
| MMAR_1123 | Rv3420c | RimI |
| MMAR_4496 | Rv0998 | mPat |
| MMAR_4889 | Rv0802c |  |
| MMAR_3205 | Rv2170 |  |
| MMAR_0332 | Rv0133 |  |
| MMAR_1341 | Rv3216 |  |
| MMAR_3740 | Rv2416c | Eis |
| MMAR_2836 | Rv3225c |  |
| MMAR_2615 |  |  |
| MMAR_0151 |  | Eis1 |
| MMAR_3692* | Rv1347c | MbtK |
| MMAR_2744 |  |  |

Several of the NATs were either established or predicted lysine acetyl transferases [KATs in Fig 1A indicated by open bars, (21, 24, 26, 50–56)], and are the focus of this study. To test if the predicted NAT impacted ESX-1 activity, we added an additional copy of the ESX-1 system to the NAT deletion strains and tested for restoration of hemolysis. We introduced the pRD1-2F9 cosmid containing the extended RD1 locus from *M. tuberculosis*, which includes the majority of the ESX-1 component and substrate genes (57), into the *M. marinum* strains lacking individual KAT genes, as well as the WT and Δ*eccCb_1_* strains. The addition of a second copy of the ESX-1 locus to the WT strain did not significantly impact hemolysis (Fig. S2B). Because the *eccCb_1MT_*(MT = *M. tuberculosis*) gene is included on the pRD1-2F9 cosmid, the pRD1-2R9 plasmid complement-ed the hemolytic activity of the Δ*eccCb_1_* strain. Interestingly, the pRD1-2F9 cosmid significantly increased the hemolytic activity of the Δ*pat* and Δ*3205* strains (*P*<0.0001, relative to the parent strains), supporting that Pat and MMAR_3205 impact hemolytic activity through the ESX-1 system. The pRD1-2F9 cosmid did not significantly impact the hemolytic activity of the Δ*rimI,* Δ*eis* or Δ*mbtK* strains, suggesting a specific connection between Pat and MMAR_3205.

We next tested if the deletion strains had altered production of PDIM, phenolic glycolipids (PGLs) and tri-acyl glycerides (TAG). We extracted total lipids from the *M. marinum* strains lacking individual KAT genes and their respective complementation strains and measured PGLs, PDIM and TAG using TLC. The WT strain, but not the Δ*mas* strain produced PDIM, TAG (Fig. 1B, top TLC) and PGL [bottom TLC, lane 1 and 2, (44, 46, 58–63)]. Both PDIM and PGL were reduced and TAG levels were elevated in total lipids extracted from the Δ*mbtK* strain (lane 3). Expression of the *mbtK* gene in the Δ*mbtK* strain increased PDIM and PGL, but did not reduce TAG levels compared to the Δ*mbtK* strain (lane 4). Deletion or complementation of the other potential KAT genes did not noticeably impact the lipids tested under these conditions (lanes 5-14). From these data, we conclude that MbtK is required for the production of WT levels of PDIM and PGLs.

### Mycobactin deficiency is linked to reduced growth and PDIM and PGL production in *M. marinum*

MbtK in is required for synthesizing mycobactin, a secreted siderophore required for iron scavenging (55). In the absence of MbtK, *M. tuberculosis* growth *in vitro* was severely impacted due to the loss of mycobactin production (55). Likewise, we found that the Δ*mbtK* strain exhibited significantly reduced growth compared to the WT and complemented strains (Fig. S3A). We next tested if the reduced growth and PDIM production in the Δ*mbtK M. marinum* strain could be rescued by supplementing the Δ*mbtK* strain with ferric Mycobactin J. As shown in Figure S3C, addition of mycobactin to the growth media restored growth of the Δ*mbtK* strain, similar to the complemented and WT strains. We grew the *M. marinum* strains in the presence of a range of mycobactin concentrations (from 2 μg/ml to 0.0002 μg/ml or in the absence of mycobactin) and extracted total lipids. Using TLC, we found that PDIM and PGL production was restored to WT levels in the presence of 2 μg/ml or 0.02 μg/ml mycobactin (Fig. S3C). 0.0002 μg/ml did not restore PDIM or PGL production to the Δ*mbtK* strain From these data we conclude that the addition of mycobactin J to the grown media transcomplemented the growth defect and PDIM/PGL production of the Δ*mbtK* strain. These data support the idea that PDIM and PGL production was reduced in the Δ*mbtK* strain due to a deficiency in iron scavenging.

MbtK, and its role in mycobactin synthesis, was previously linked to phospholipid homeostasis (55). However, MbtK has not previously been linked to PDIM and PGL production. To gain insight into the mechanisms underlying the phenotypes associated with the loss of MbtK we used proteomics to measure the changes in protein levels. We generated cell-associated and secreted protein fractions from the WT, Δ*mbtK,* and the Δ*mbtK*/p*mbtK* strains (Dataset S1) and measured protein levels in each fraction using nUHPLC MS-MS/MS quantitative bottom-up proteomics (44, 64). The deletion of *mbtK* resulted in a significant increase in 15 proteins (Figure S4A), the majority of which are associated with three chromosomal regions associated with iron uptake, including ESX-3 (EccB3 and MycP3), MMAR_2874, MMAR_2875, cyp278A1 and MMAR_2878, which are colocalized on the chromosome, and MbtF, MbtE and MbtD, which are colocalized with MbtK on the chromosome. The Cyp278A1 protein is annotated as being involved in heme uptake (65). The additional genes are conserved hypothetical proteins. Expression of *mbtK* in the Δ*mbtK* strain restored protein levels to those in the WT strain. We also measured the protein levels in the secreted fractions. The levels of the Mbt proteins were increased in the secreted fraction from the Δ*mbtK* strain as compared to the strains expressing *mbtK*. There was no significant change in the secretion of any ESX proteins to explain the reduced hemolytic activity in Fig. 1A. These data did not allow us to connect the loss of MbtK to the production of PDIM and PGL.

### Overexpression of *eis* restores virulence lipids to the Δ*mbtK* strain

To understand the requirement for MbtK in the lipid biosynthesis, we asked if constitutive expression of the other putative KAT genes could restore wild-type PDIM, PGL and TAG levels to the Δ*mbtK* strain. We introduced each KAT gene into the Δ*mbtK* strain on an integrating plasmid, expressed from the MOP promoter, and confirmed the presence of the plasmid in each strain by PCR (Fig. S5). The expression of the *rimI, pat* or *MMAR_3205* genes did not detectably impact the levels of PDIM, TAG or PGLs in the Δ*mbtK* strain (Fig. 2A, lanes 5-7). However, constitutive expression of the *eis* gene specifically restored PDIM, TAG and PGL levels in the Δ*mbtK* strain (lane 8) to those in the WT and complemented strains.

**Figure 2.**
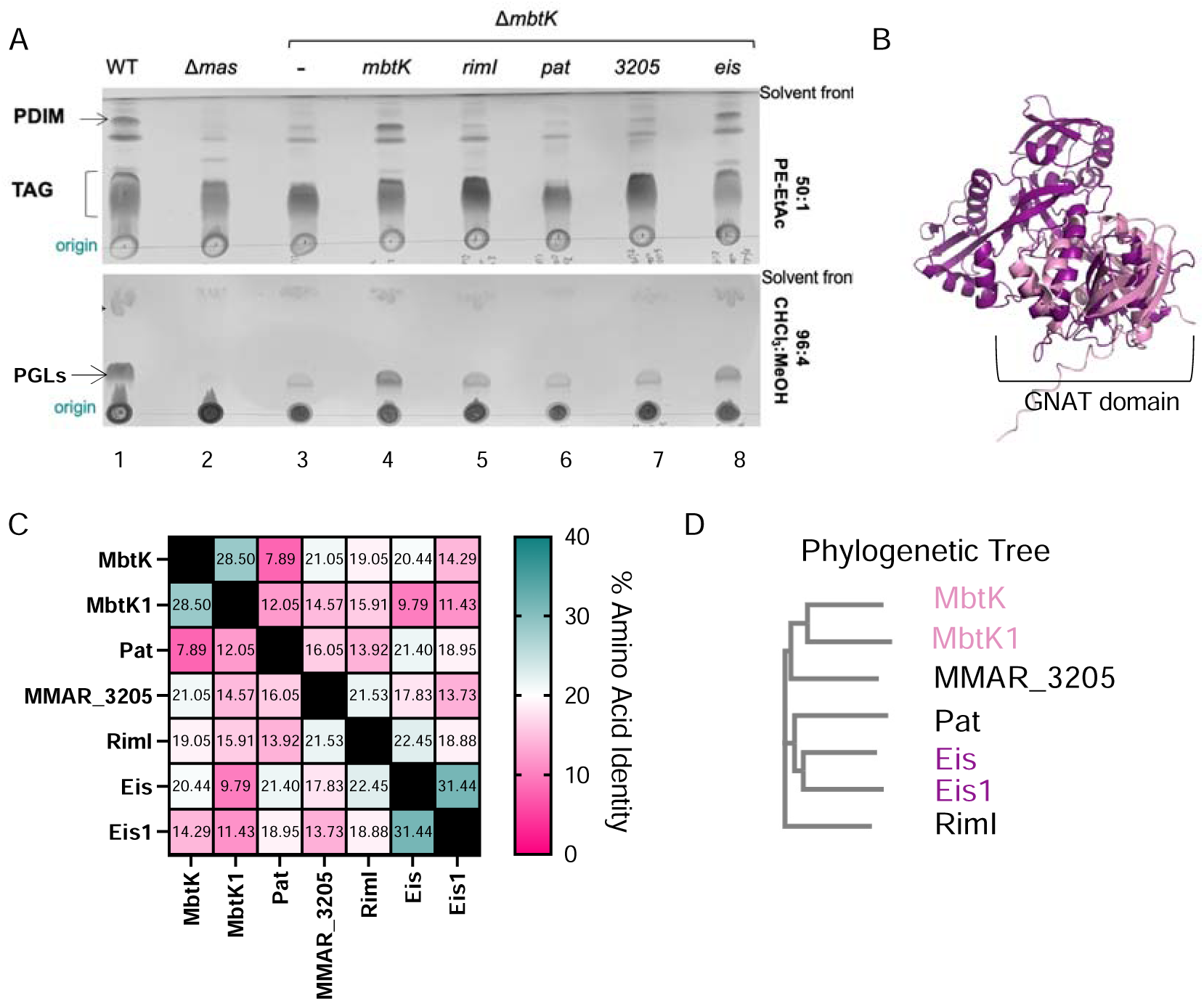
Overexpression of Eis functionally restores PDIM production in the Δ*mbtK* strain. **A.** TLC analysis of total lipids from the indicated *M. marinum* strains. The NAT genes in lanes 4-8 were constitutively expressed in the ΔmbtK strain using the mycobacterial optimal strain. 6 µl of total lipids was separated for each strain. The indicated solvents and lipids visualized are indicated. The TLC is representative of at least three biological replicates. **B.** Overlay of the predicted structures for MbtK and Eis NATs. The PDB files used were: https://alphafold.ebi.ac.uk/files/AF-B2HMD6-F1-model_v3.pdb and https://alphafold.ebi.ac.uk/files/AF-B2HM88-F1-model_v3.pdb. The structures were overlaid using PyMOL v.2.4.0. **C**. % amino acid identify of the NATs from panel A. Protein sequences were obtained from Mycobrowser (65), and aligned using Clustal Omega (95). **D.** Clustering by phylogenetic neighbor joining tree without distance corrections, generated based on protein alignments generated by Clustal Omega.

Eis is a N-acetyltransferase that is clinically linked to aminoglycoside resistance in *M. tuberculosis* (28, 66–71). To our knowledge, there are no known connections between the Eis and MbtK acetyltransferases. We first considered if Eis restored lipid production simply because it was the most similar NAT to MbtK. Based on structural predictions, MbtK and Eis proteins do not share domains beyond the conserved GNAT domains (AlphaFold, Fig 2B). The RimI, Pat or MMAR_3205 proteins that did not functionally complement the Δ*mbtK* also share the GNAT domain. At the amino acid level, Eis, RimI and MMAR_3205 are all ∼20% identical to MbtK (Fig. 2C), and both MbtK1 and MMAR_3205 are more phylogenetically related KAT to MbtK the Eis (Fig. 2D). Therefore, we hypothesized that Eis specifically restored PDIM/PGL production to the Δ*mbtK* strain. Our proteomics analysis shows that expression of *eis* in the Δ*mbtK* strain restored the levels of 12 of the 15 previously discussed proteins, with the exception of MMAR_2874, MMAR_2875 and Cyp278A1, which are at the same genetic locus (Fig. S4). These data suggest that *eis* partially complements the Δ*mbtK* strain.

*M. tuberculosis* required supplementation with iron during laboratory culture when the *mbtK_MT_* gene was deleted (55). The Δ*mbtK* (*MMAR_3692*) *M. marinum* strain exhibited reduced growth without the addition of supplemental iron compared to the WT strain (Fig. S3A). An orthologous *mbtK* gene, *MMAR_1587*, in *M. marinum* genome which may allow growth of the Δ*mbtK* strain without additional iron. MMAR_1587 is 28.5% identical at the amino acid level, and is predicted to be structurally similar to MbtK (Fig. S6A) MMAR_1587 is 54.73% identical to the MbtK*_MT_*protein at the amino acid level, and is annotated as the MbtK ortholog on Mycobrowser (15). Although the MMAR_3692 protein is 26% identical to MbtK*_MT_,* the *MMAR_3692* gene is syntenic with the *mbtK_MT_.* As such, we refer to *MMAR_3692* as *mbtK,* and *MMAR_1587* as *mbtK1.* (Fig. 2D). *M. marinum* has an orthologous *eis* gene, *MMAR_0151,* which is annotated as *eis1* (65). The Eis and Eis1 proteins are 31.44% identical, and are predicted to be structurally similar (Fig. S6A). To further probe the interaction between MbtK and Eis, we constitutively expressed *mbtK1* and *eis1* in the Δ*mbtK* strain. Constitutive expression of the *mbtK1* or *eis1* genes did not completely rescue PDIM or PGL levels in the Δ*mbtK* strain as visualized by TLC (Fig. S6B). These data suggest that additional copies of MbtK or Eis are functionally redundant, and that Eis likely specifically rescues PDIM, PGL and TAG production in the Δ*mbtK* strain.

### Eis N-terminally acetylates PapA5 *in vitro*

The proteomics data did not provide an obvious mechanism for how Eis restored PDIM and PGL production to the Δ*mbtK* strain. We demonstrated that several proteins required for PDIM and/ or PGL production (PpsA, PapA5, Mas and PpsE, Pks15/1) were N-terminally acetylated [Fig S7, Pink, (*1*)]. Although neither MbtK nor Eis have reported to N-terminally acetylate proteins, we tested if MbtK and/or Eis could N-terminally acetylate one or more biosynthetic proteins required for PDIM and PGL production.

It was previously established that the first six amino acids of a substrate protein were sufficient for N-terminal acetylation *in vitro* (30, 72). We had N-terminal six AA peptides synthesized based on the N-terminal peptides we previously measured using proteomics [Fig. 3A, (16)]. Only the Mas peptide contained a “K” residue, suggesting that we could measure N-terminal acetylation of these peptides. The N-terminal peptides were used as substrates for *in vitro* acetylation assays with the WT or inactive versions of MbtK (MbtKE161A, active site mutation as predicted by UniProt) and Eis [EisY128A, proton donor active site mutant based on (29)] that we expressed and purified from *E. coli* (Fig. S8). Transferring the acetyl group from Acetyl-CoA to a peptide substrate results in a free thiol group on the resulting CoA molecule. The reaction of this free thiol with 5, 5’-dithiobis-(2-nitrobenzoic acid) DTNB results in a colorimetric change that can be measured at 412nm. The resulting product has 1:1 stoichiometry with the acetylated peptide and can be used to quantify protein acetylation (29, 73). The RimI NAT from *Salmonella* with the S18 substrate [(72, 74), Fig. S8] was a positive control for the reaction. Incubation of the PpsE (*P*=0.0147) or PapA5 (*P*<0.0001) peptides with the WT MbtK resulted in significantly higher DTNB production than incubation with the MbtKE161A protein (Fig. 3B). Incubation of the PpsA, Mas or Pks15/1 peptides resulted in product formation that was not significantly different between the active and inactive enzymes. These data support that MbtK may acetylate the N-terminal amino acid of PpsE or PapA5.

**Figure 3.**
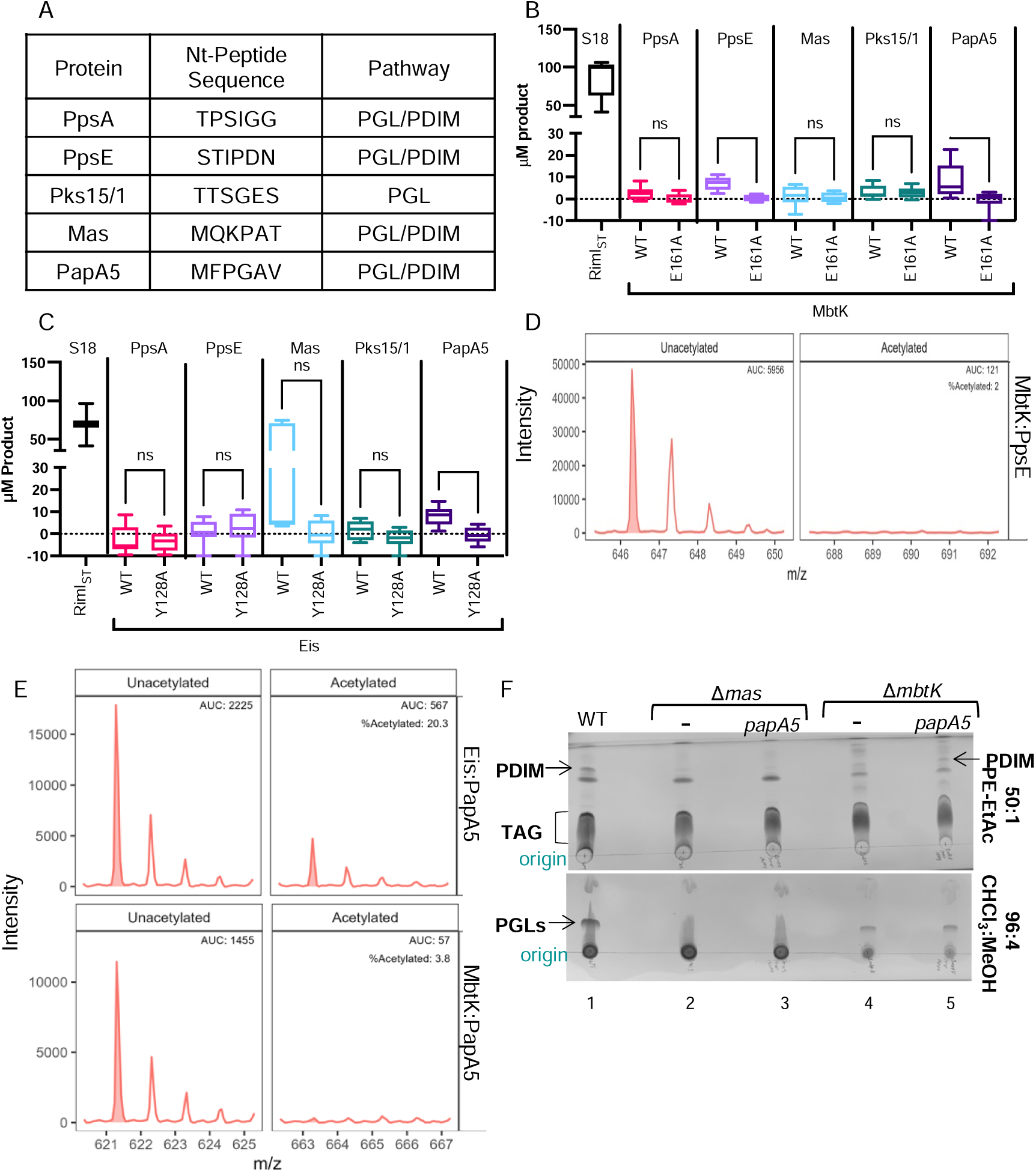
PapA5 is N-terminally acetylated by Eis *in vitro.* **A.** Table of peptides from proteins in the PDIM/PGL pathways. B. DTNB assay measuring the production of TNB (412nm) upon the acetylation of the PpsA, PpsE, Mas, Pks15/1 and PapA5 peptides by **B.** WT or inactive MbtK and **C.** the RimI protein from *Salmonella typhimurium* with the S18 N-terminal peptide (ARYFRR). DTNB assays were performed in technical triplicate. The data represent the mean product formation from all three experiments. Significance was determined using a Student’s t-test between the active and inactive forms of each NAT protein. B.* *P*=0.0417, **** *P*<0.0001 C. ** *P*=0.0096**. D.** Flow injection analysis of the N-terminal peptide of PpsE with purified MbtK protein. Left panel: peaks corresponding to the unacetylated N-terminal peptide of PpsE. Right panel: mass range corre-sponding to the acetylated version. **E.** Flow injection of the N-terminal peptide of PapA5 incubated with either purified Eis protein or purified MbtK protein. Top left panel: peaks corresponding to unacetylated PapA5. Top right: the acetylated version of PapA5 when incubated with Eis. Bottom left panel: peaks corresponding to unacetylated PapA5, Bottom right panel: the same mass range corresponding to the acetylated version of PapA5. **F.** TLC on total lipids from *M. marinum.* 6μl total lipid loaded into each lane, PDIM run 3x consecutively in the solvent indicated, PGLs run once in the solvent indicated. PDIM lipids visualized by phosphomolybdic acid and subsequent charring, PGL lipids visualized by 1% a-naphthol and subsequent charring. The TLC is representative of 3 biological replicates, the remaining 2 are shown in Fig. S9.

We conducted the same experiment using Eis or EisY128A, a proton donor active site mutant (29). Incubation of the PapA5 peptide with the WT Eis protein resulted in significantly higher DTNB production than incubation with the Eis Y128A protein *(P*=0.0096, Fig. 3C). None of the other peptides tested exhibited significantly reduced product formation between the WT and dead enzymes. These data are consistent with the possibility that Eis may N-terminally acetylate PapA5.

To directly test the acetylation of PpsE and PapA5 by MbtK and Eis, we used FIA mass spectrometry. As shown in Figure 3D, MbtK did not N-terminally acetylate the PpsE peptide. Although we detected the unacetylated peptide (left), we did not detect the acetylated peptide. We likewise tested if Eis or MbtK could N-terminally acetylate the PapA5 peptide. The unacetylated PapA5 peptide was detected when mixed with either Eis (Fig. 3E, top left) or MbtK (bottom left). The acetylated PapA5 peptide was detected in the presence of Eis, but not MbtK (Fig. 3E, right). We conclude that Eis, but not MbtK, N-terminally acetylates PapA5 *in vitro*. These data suggest that *eis* expression in the Δ*mbtK* strain restores PDIM production by N-terminal acetylation of the PapA5 protein (Fig. 3F). We attempted to measure N-terminal acetylation in the Δ*mbtK* strain, and in the corresponding complementation and *eis* expression strain (Dataset S1). We were unable to measure the N-terminal peptide of PapA5 in these strains. We did not identify N-terminal peptides whose acetylation in the Δ*mbtK* strain was significantly decreased and was restored by both the *mbtK* and *eis* expression, or by expression of the *eis* gene. Potential candidates for targets that did not reach significance are in (Fig. S4B). These data did not allow us to determine if the N-terminal acetylation of PapA5 was promoted by *eis* expression in *M. marinum*.

We tested if *papA5* could restore PDIM/PGL production or growth to the Δ*mbtK* strain by collecting total lipids and visualizing PDIM and PGL on TLCs. Expression of *papA5* did not increase PDIM or PGL Levels (Fig. 3F, lanes 2 and 3). Expression of the *papA5* gene in the Δ*mbtK* strain increased the levels of PDIM and PGLs in this strain, but not to WT levels (lanes 4 and 5, and additional replicates in Fig S9). Together these data support the conclusion that although PapA5 partially rescued the reduced PDIM/PGL levels in the Δ*mbtK* strain, it did not rescue growth under the conditions tested.

### The requirement of MbtK for mycobacterial pathogenesis can be suppressed by Eis or PapA5

Because the Δ*mbtK_MT_*was attenuated for virulence in a mouse model of infection (55), we tested if the Δ*mbtK* strain was attenuated in two models of mycobacterial infection. First, we tested if *mbtK* was required for macrophage cytolysis using Ethidium homodimer (Eth-D1), which is a membrane impermeable nucleic acid stain (*16*). If *M. marinum* infection perforates the macrophage membrane, the cells stain red. We infected RAW 264.7 cells with *M. marinum* at an M OI of 5, and counted the presence of red cells (Fig. 4A). Infection of RAW 264.7 cells with wild-type *M. marinum* (*P*<0.0001), but not the Δ*eccCb_1_* strain, resulted in significant levels of cytolysis compared to the uninfected cells. The Δ*mas* strain and the Δ*mbtK* strains resulted in significantly less cytolysis than the WT strain (*P*<0.0001). The complementation strains significantly increased cytolysis compared to the deletion strains (*P*<0.0001) but were still significantly lower than the WT strain (*P*<0.0001and *P*=0.0090, respectively). The Δ*mbtK* strain expressing the *mbtKE161A* gene did not significantly affect cytolysis compared to the Δ*mbtK* strain. From these data we conclude that the NAT activity of MbtK is required for macrophage cytolysis.

**Figure 4.**
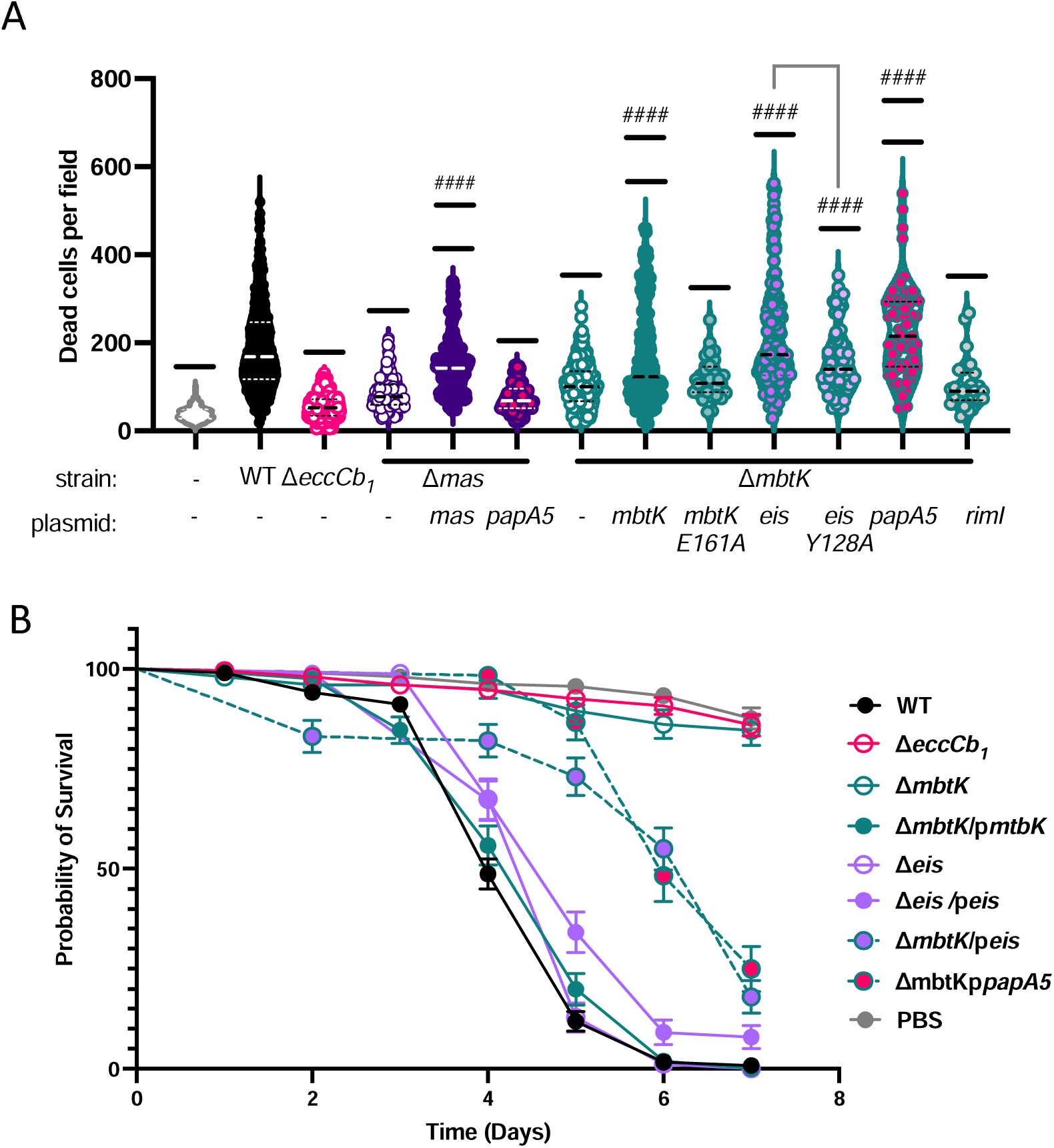
The requirement of MbtK for mycobacterial pathogenesis can be suppressed by constitutive expression of Eis or PapA5. **A**. RAW 264.7 cells infected with *M. marinum* strains at an MOI of 5. The data is the average of at least 3 independent infections, each performed in biological triplicate. **** *P*<0.0001, *** *P*=0.0001, ** *P*=0.0218, #### *P*<0.0001. **B.** *Galleria mellonella* infected with the *M. marinum* strains indicated. Larvae were injected with 1×10^7^ bacteria into the right hindmost proleg. Larvae were infected in 3 groups of 10, for a total of 30 larvae per strain per infection. Data is representative of at least 3 biological replicates, each performed in technical triplicate. Statistical analysis was performed as in (79). Survival curves were compared using a log-rank Mantel-Cox test, *P*<0.0001.

We tested macrophage cytolysis following infection with the Δ*mbtK* strain constitutively expressing the *eis* or the *eisY128A* gene (Fig. 4B). Expression of the *eis* or *eisY128A* gene in the Δ*mbtK* strain significantly increased cytolysis compared to the Δ*mbtK* strain (*P*<0.0001), similar to the WT strain. Infection with the Δ*mbtK* strain expressing the *eisY128A* gene resulted in significantly reduced cytolysis compared to the *eis* gene (*P*=0.0001). We conclude that constitutive expression of *eis* suppresses the loss of MbtK during macrophage infection, and that this suppression requires the NAT activity of Eis.

We tested if constitutive expression of *papA5* would likewise restore cytolysis to the Δ*mbtK* strain. Infection of RAW 264.7 cells with the Δ*mbtK* strain constitutively expressing *papA5* resulted in significantly more cytolysis than the WT (*P*=0.0218) or Δ*mbtK* strains (*P*<0.0001, Fig. 4A). To test if suppression of cytolysis by *papA5* expression was specific we added two controls. First, constitutive expression of *papA5* in the Δ*mas* strain failed to restore cytolysis to the Δ*mas* strain. Second, constitutive expression of the *rimI* NAT in the Δ*mbtK* strain failed to restore cytolysis to the Δ*mbtK* strain. Collectively, our data support that acetylation of PapA5 by Eis bypasses the loss of MbtK.

We tested if the orthologous *mbtK1* and *eis1* genes were essential for macrophage cytolysis. Expression of the *mbtK1* gene partially restored cytolysis to the Δ*mbtK* strain (Fig. S7B) Expression of the *eis1* gene in the Δ*mbtK* strain resulted in cytolysis levels significantly higher than the WT strain (*P*<0.0001). Deletion of the *mbtK1* or *eis* genes significantly reduced macrophage cytolysis compared to the WT strain (*P*<0.0001), but cytolysis was not restored upon expression of *mbtK1* or *eis* in these strains. Deletion of the *eis1* gene resulted in a significant reduction of macrophage cytolysis compared to the WT strain, and this was significantly increased upon expression of the *eis1* gene, though not to WT levels.

To complement the cellular model of infection, we tested the ability of *M. marinum* to kill the waxworm, *Galleria mellonella* larvae*. G. mellonella* was previously established as a model for mycobacterial infection (75–79). For each experiment, we infected 30 worms, each injected with 1×10^7^ bacteria in a 5 μL volume. Experiments were performed in biological triplicate. As shown in Figure 4B, WT *M. marinum* killed all of the infected *G. mellonella* by 7 days post-infection. In contrast, infection with either the Δ*eccCb_1_* or Δ*mbtK* strains resulted in less killing, similar to the uninfected worms. Deletion of *eis* did not significantly impact the killing of *G. mellonella.* Expression of *mbtK* restored the ability of the Δ*mbtK* strain to kill *G. mellonella,* similar to the WT strain. Expression of either *eis* or *papA5* in the Δ*mbtK* strain partially restored the ability of *M. marinum* to kill *G. mellonella.* Introduction of the pRD1-2F9 plasmid to the Δ*mbtK* strain or growing the Δ*mbtK* strain in ferric mycobactin did not restore *G. mellonella* killing to the ΔmbtK strain (Fig. S3B). Together, these data support that the expression of Eis or PapA5 suppresses the loss of *mbtK* in *M. marinum*.

## Discussion

We sought to identify conserved GNAT containing proteins important for the ESX-1 and PDIM virulence pathways. We found that the majority of GNAT proteins previously annotated as lysine acetyltransferases were required for wild-type levels of hemolysis by *M. marinum.* We identified several potential NATs required for hemolysis that could be rescued by the addition of a second ESX-1 locus. These GNAT proteins will be followed up in future studies. Our data support a model where mycobactin-mediated iron uptake promotes the PapA5-mediated biosynthesis of PDIM and PGLs (80). In the absence of MbtK, the production of PDIM and PGLs is reduced compared to the WT *M. marinum* strain. PDIM and PGL production was complemented by expression of either *mbtK* or *in trans* by the addition of Mycobactin J to the growth media. Complementation was dependent upon the acetyltransferase activity of MbtK, as expression of the *mbtKE161A* allele, which does not have NAT activity, did not restore PDIM and PGL production. We showed that the expression of *papA5* or the *eis* NAT in the Δ*mbtK* strain specifically suppressed the requirement of MbtK in PDIM and PGL production. The simplest mechanism, which is supported by our *in vitro* assays, is that Eis N-terminally acetylates PapA5, promoting the production of PDIM and PGL in the absence of MbtK. Overall, our studies support a previously undiscovered link between MbtK, iron uptake and the final step of PDIM and PGL biosynthesis (Fig. 5).

**Figure 5.**
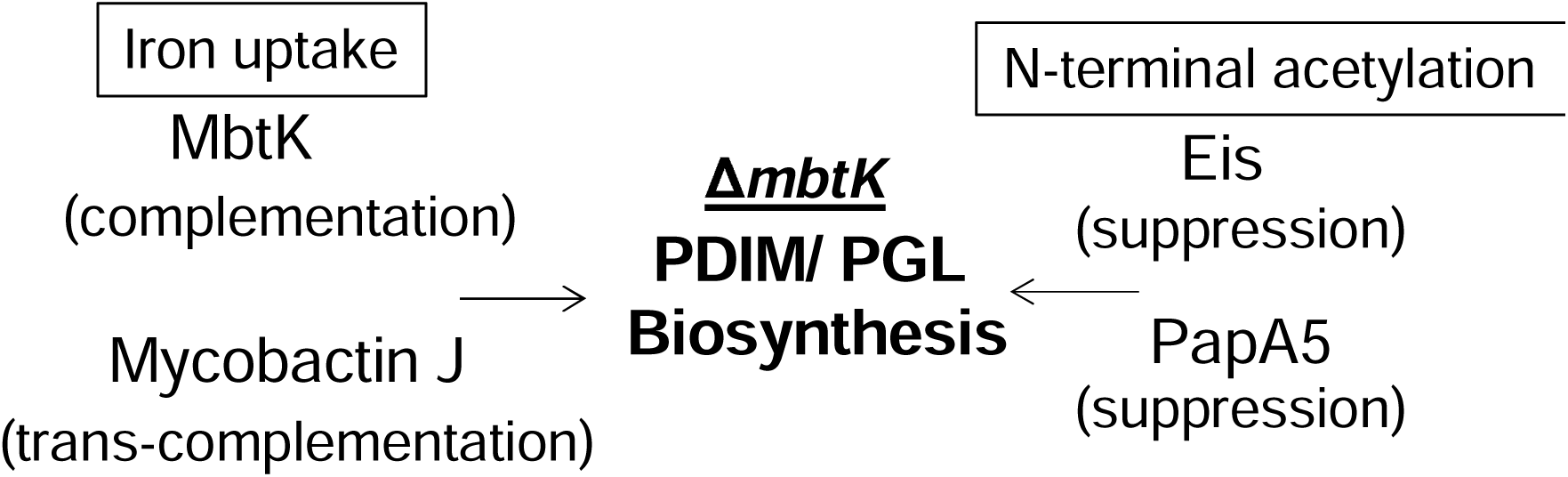
Model of Interactions between MbtK, Eis and PDIM biosynthesis. This study revealed that MbtK, likely through siderophore production, is required for the production of PDIM and PGL in *M. marinum.* This requirement was complemented in trans by the addition of Mycobactin J. Constitutive expression of either Eis, or PapA5 specifically suppressed the loss of PDIM/PGL production in the absence of MbtK, suggesting that the N-terminal acetylation of PapA5 could promote the restoration of PDIM/PGL synthesis in the Δ*mbtK* strain.

Our model is supported by several lines of evidence previously published in the literature. It was previously established by Madigan et al that MbtK is required broadly for lipid and mycobactin production (55). MbtK is required for the synthesis of the lipid tail of the mycobactin siderophore, but more broadly, the loss of MbtK and iron uptake in *M. tuberculosis* resulted in a depletion of phospholipids during iron starvation (55). Although Madigan et al connected lipid production to MbtK mediated siderophore iron up-take in *M. tuberculosis,* the PDIM and PGL lipids were not readily measured using lipidomics approaches in this study. As such, the connection between mycobactin dependent iron uptake and PDIM/PGL biosynthesis has not been previously investigated. The connection between iron and PDIM/PGL biosynthesis could be direct. For example, the PapA5 mediated synthesis step in PDIM biosynthesis could require iron. In support of this, *papA5* overexpression or the addition of mycobactin restored PDIM/PGL biosynthesis in the Δ*mbtK* strain. Alternatively, the connection between iron and PDIM/PGL biosynthesis could be indirect. For example, it is possible that iron levels are a signal that regulate the production of PDIM and PGL. Further experimentation is required to differentiate between these two possibilities.

The mechanisms connecting MbtK to mycobacterial pathogenesis are unclear. Yet, it was surprising that constitutive expression of the *papA5* or *eis* genes fully restored macrophage cytolysis to the Δ*mbtK* strain. Infection of *G. mellonella* revealed that increasing PapA5 or Eis restored virulence to the Δ*mbtK* strain, but the kinetics of killing differed from that of the WT strain. These data support that perhaps expression of these genes restored PDIM/PGL production, but did not fully complement the Δ*mbtK* strain. Further work is required to determine if constitutive expression of *eis* or *papA5* restore more that the PDIM/PGL deficiency to the Δ*mbtK* strain.

In this study, we uncovered a link between Eis and PapA5. Eis is a bifunctional N-acetyl-transferase that acetylates aminoglycoside antibiotics to inactivate them. Mutations that lead to *eis* overexpression naturally occur in clinical isolates of *M. tuberculosis,* leading to kanamycin resistance (28, 66–71). Numerous studies have reported inhibitors of Eis as a potential path to prevent aminoglycoside resistance during *M. tuberculosis* infection (81–90). Eis reversibly acetylates lysine residues in the HupB protein, affecting chromosomal condensation in *M. tuberculosis* (26). We showed that Eis N-terminally acetylates PapA5 *in vitro*, adding another potential activity to this enzyme. We were unable to detect the N-terminal peptide of PapA5 in our proteomics experiment. However, expression of *papA5* suppressed several phenotypes of the Δ*mbtK* strain supporting this connection (Fig. 5). If Eis acetylates the N-terminus of PapA5 *in vivo*, this modification must promote PapA5 activity, allowing suppression of the PDIM/PGL defect in the Δ*mbtK* strain. Our data further support an emerging literature that NATs, including Eis, are multifunctional (91).

Knobloch et al previously suggested that the orphan *mbtK* gene (*MMAR_1587*) was the *M. marinum* ortholog of *M. tuberculosis* MbtK (92). Despite being located at the *mbt* locus, *MMAR_3692* was not mentioned in this publication. In our study, we found that deletion of the *MMAR_3692* gene resulted in reduced growth during standard culturing conditions. The growth defect was significant, but unlike in *M. tuberculosis,* the deletion of *mbtK* did not abrogate growth. Similar to *M. tuberculosis,* supplementation with mycobactin restored growth of the Δ*mbtK* strain (55). Deletion of the *mbtK* gene, but not the *MMAR_1587* gene resulted in a measurable PDIM/PGL defect. Likewise, expression of *MMAR_1587* in the Δ*mbtK* strain did not restore PDIM levels. From our studies, we propose that the *MMAR_3692* gene is the *M. marinum* ortholog of the *M. tuberculosis mbtK*. Our data also suggests that the two genes in *M. marinum* are functionally distinct.

In non-tubercular mycobacterial species, there are two functionally distinct eis genes. In *M. abscessus,* these are annotated as Eis1Mab and Eis2mab (93). Overexpression of *eis2mab,* but not *eis1mab,* resulted in resistance to Kanamycin. We likewise found that there are two potential *eis* genes in *M. marinum.* Deletion of either *eis* gene attenuated *M. marinum* in a macrophage model of infection (Fig. S6C). Overexpression of *eis,* but not *eis1* in *M. marinum* resulted in low level Kan resistance (Fig S10), supporting that robust literature linking Eis levels to aminoglycoside resistance, and that the two *eis* genes are functionally distinct. Our macrophage infection data supports that overexpression of *eis1* restored cytolysis of the Δ*mbtK* strain to greater than WT levels (Fig. S6). Yet, expression of *eis1* did not restore PDIM/PGL production to the Δ*mbtK* strain. The mechanisms connecting Eis1 to MbtK need further experimentation.

Overall, this study revealed a new connection between iron scavenging and PDIM and PGL biosynthesis that requires further investigation. We uncovered that the Eis NAT, which has been widely linked to drug resistance in *Mycobacterium,* also has the potential to N-terminally acetylate proteins, including those required for the production of the PDIM and PGL lipids. Our data support that NATs cannot be characterized by the targets they modify, and likely play unexpected, interconnected and as of yet undefined roles in mycobacterial pathogenesis and physiology.

## Materials and Methods

A summary of the methods are found here. Where necessary, detailed methods are provided in the Supplementary Information. *Mycobacterium marinum* strains were derived from the *Mycobacterium marinum* M strain (ATCC BAA-535), and maintained as described previously. In frame deletion strains were generated using allelic exchange, and can be found in TableS1. Complementation plasmids, found in Table S2, were constructed using the primers in Table S3 using FastCloning. Site directed mutagenesis was performed as described previously (94). Hemolysis assays were performed as described. RAW 264.7 cells were maintained as described and infected with *M. marinum* at an MOI of 5. Macrophage cytotoxicity were measured by measuring Ethidium homodimer uptake as described. *G. mellonella* larval infections were performed by in-jecting *M. marinum* at a concentration1×10^7^/5µL. Thin-layer chromatography experiments was performed as described in (46). Expression and purification of 6xHis-tagged proteins were expressed in *E. coli,* and purified using Ni-NTA resin, followed by removal of the epitope tag using TEV. Proteins were used to test activity in vitro against N-terminal peptides (Genscript) using <u>5,5-dithio-bis-(2-nitrobenzoic acid) (DTNB) assays as</u> (73) with minor modifications. Peptide acetylation following the DTNB assay was confirmed using MALDI. Bottom-up proteomics was performed after labelling samples with deuterated acetyl groups as (16) to quantify protein levels and N-terminal acetylation. Label-free quantification (LFQ) was used for differential protein expression analysis, and data were processed using R (v4.3.2). The code can be found on github (https://github.com/Champion-Lab/3692). Rawdata is available through MassIVE and The Proteome Exchange at ftp://MSV000095200@massive.ucsd.edu with Identifier MSV000095200 PXD053528.

## Supporting information

Proteomics dataset

Supplementary Figures Tables Methods and References

## Acknowledgments

We would like to thank Bill Boggess at the Notre Dame Mass Spectrometry and Proteomics Facility for continued assistance. Research reported in this publication was supported by NIAID of the National Institutes of Health under award number AI106872. BSJ was supported by a Graduate Fellowship from the Eck Institute for Global Health at the University of Notre Dame. We thank Dr. Micah Ferrell, who constructed several NAT deletion strains used in Figure S1.

