## Supplementary Figures Tables Methods and References for "N*-*acetyl-transferases required for iron uptake and aminoglycoside resistance promote virulence lipid production in *M. marinum*"

**This File Includes:**

**Supplementary Figures**

**Figure S1: Confirmation of *Mycobacterium marinum* deletion and complementation strains.**
**Figure S2: Hemolytic activity of *M. marinum* NAT deletion strains**
**Figure S3: Ferric mycobactin J rescues growth and PDIM/PGL production of, but not *G. mellonella* killing by the Δ*mbtK* strain**
**Figure S4. Proteomic measurement of cell associated and secreted protein levels and percent N-terminal acetylation of *M. marinum* proteins.**
**Figure S5. *Mycobacterium marinum* cross-complement strain confirmations.**
**Figure S6: Cross-complementation phenotypes of Δ*mbtK* with *mbtK1* and *eis1*.**
**Figure S7: PDIM and PGL biosynthetic pathway.**
**Figure S8: Heterologous expression and purification of NAT**
**Figure S9: PapA5 restores PDIM/PGL to the Δ*mbtK* strain.**
**Figure S10: Kanamycin resistance capacity of strains constitutively expressing *eis*.**

**Supplementary Tables**

**Table S1**: List of Strains
**Table S2:** List of Plasmids
**Table S3:** List of Primers

**Dataset S1**

**Supplementary Methods**

**Supplementary References**

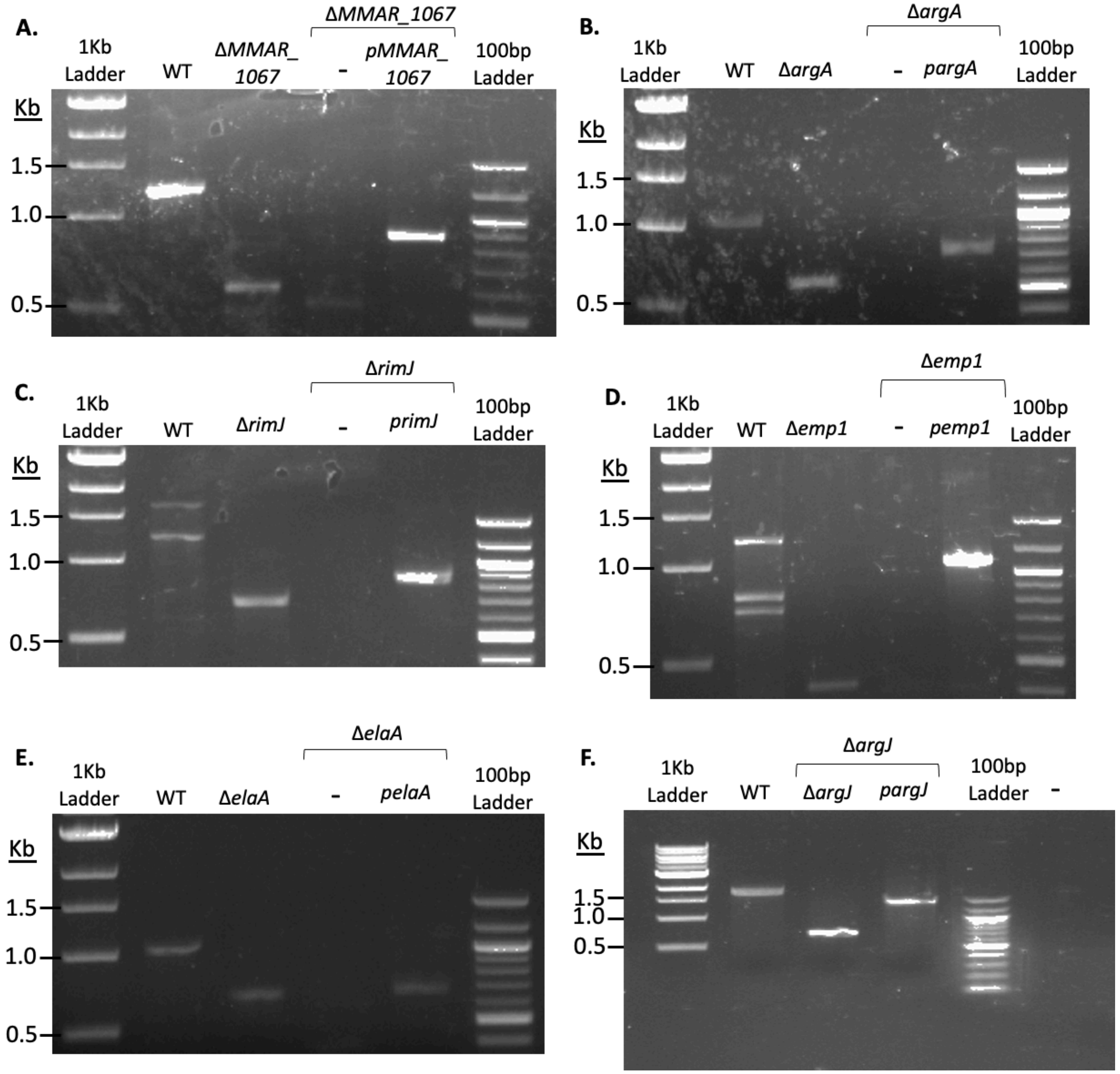


**Figure S1: Confirmation of *Mycobacterium marinum* Deletion and Complementation Strains:** PCR confirming deletion and complementation of **A.** *MMAR_1067* gene. Product: WT 1,222 bp, Δ*MMAR_1067* 592 bp. Size plasmid product: 876 bp. **B.** *argA* gene. Product: WT 1,051 bp, Δ*argA* 593 bp. Size plasmid product: 735 bp. **C.** *rimJ* gene. Product: WT 1,233 bp, Δ*rimJ*: 687 bp. Size plasmid product: 864 bp. **D.** *emp1* gene. Product: WT 1,219 bp, Δ*emp1*, 364 bp. Size plasmid product: 1,065 bp. **E.** *elaA* gene. Product: WT: 1,036 bp. Δ*elaA*: 670 bp. Size plasmid product: 681 bp. **F.** *argJ* gene. Product: WT 1,798 bp, Δ*argJ*: 691 bp. Size plasmid product: 1,430 bp. PCR primers were designed based on sequence annotations from Mycobrowser (1).

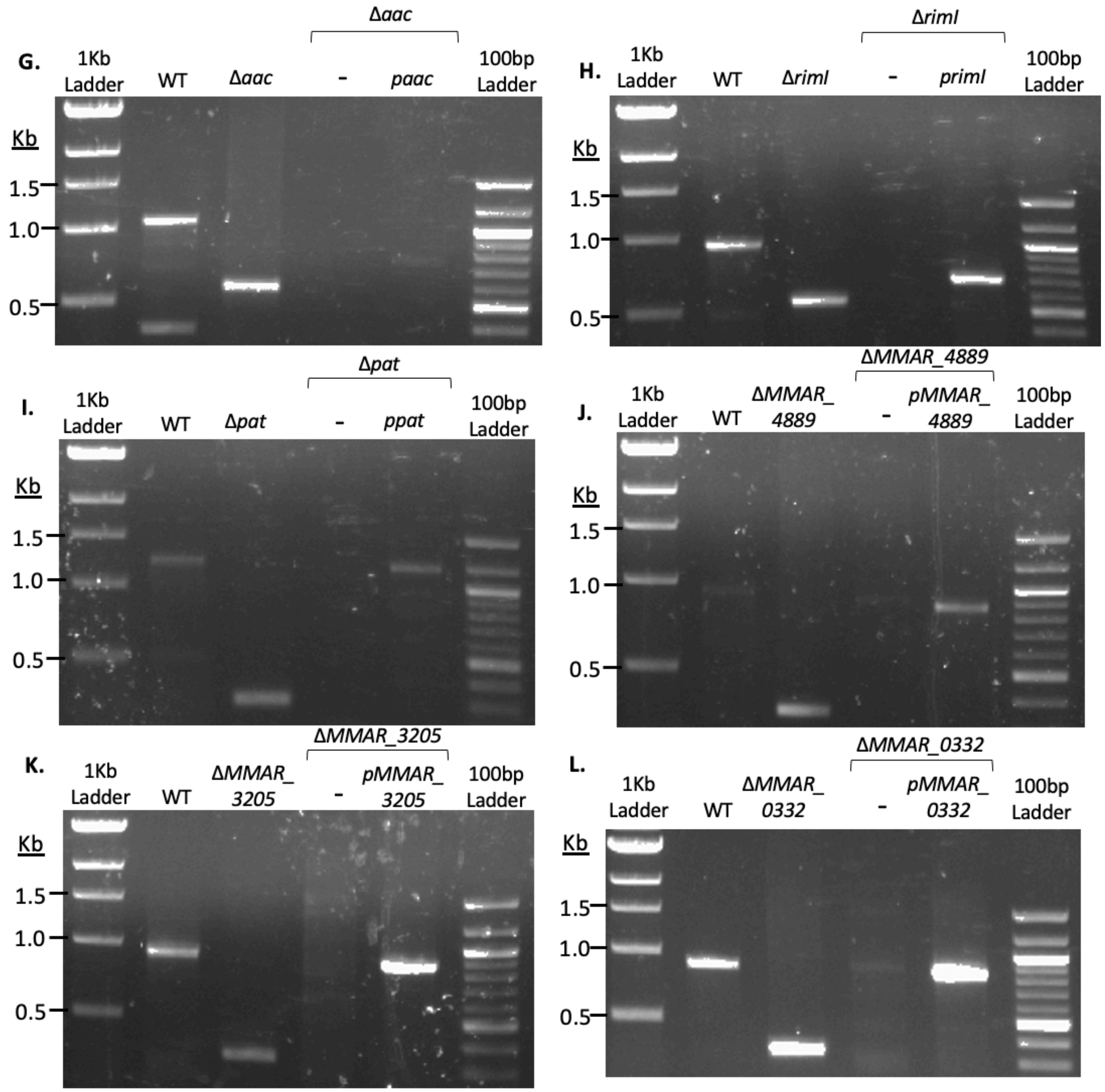


**Figure S1 (Cont.): *Mycobacterium marinum* Deletion and Complement Strain Confirmations:** PCRs confirming deletion and complementation of the **G.** *aac* gene. Product WT: 1,054 bp. Δ*aac*: 574 bp. Size plasmid product: 756 bp. Absence in control lane. **H.** *rimI* gene. Product WT: 939 bp. Δ*rimI*: 546 bp. Size plasmid product: 687 bp. Absence in control lane. **I.** *pat* gene. Product WT: 1,246 bp. Δ*pat*: 273 bp. Size plasmid product: 1,161 bp. Absence in control lane. **J.** *MMAR_4889* gene. Product WT: 934 bp. Δ*MMAR_4889*: 331 bp. Size plasmid product: 861 bp. Absence in control lane. **K.** *MMAR_3205* gene. Product WT: 924 bp. Δ*MMAR_3205*: 337 bp. Size plasmid product: 849 bp. Absence in control lane. **L.** *MMAR_0332* gene. Product in WT: 877 bp. Δ*MMAR_0332*: 337 bp. Size plasmid product: 816 bp. Absence in control lane.

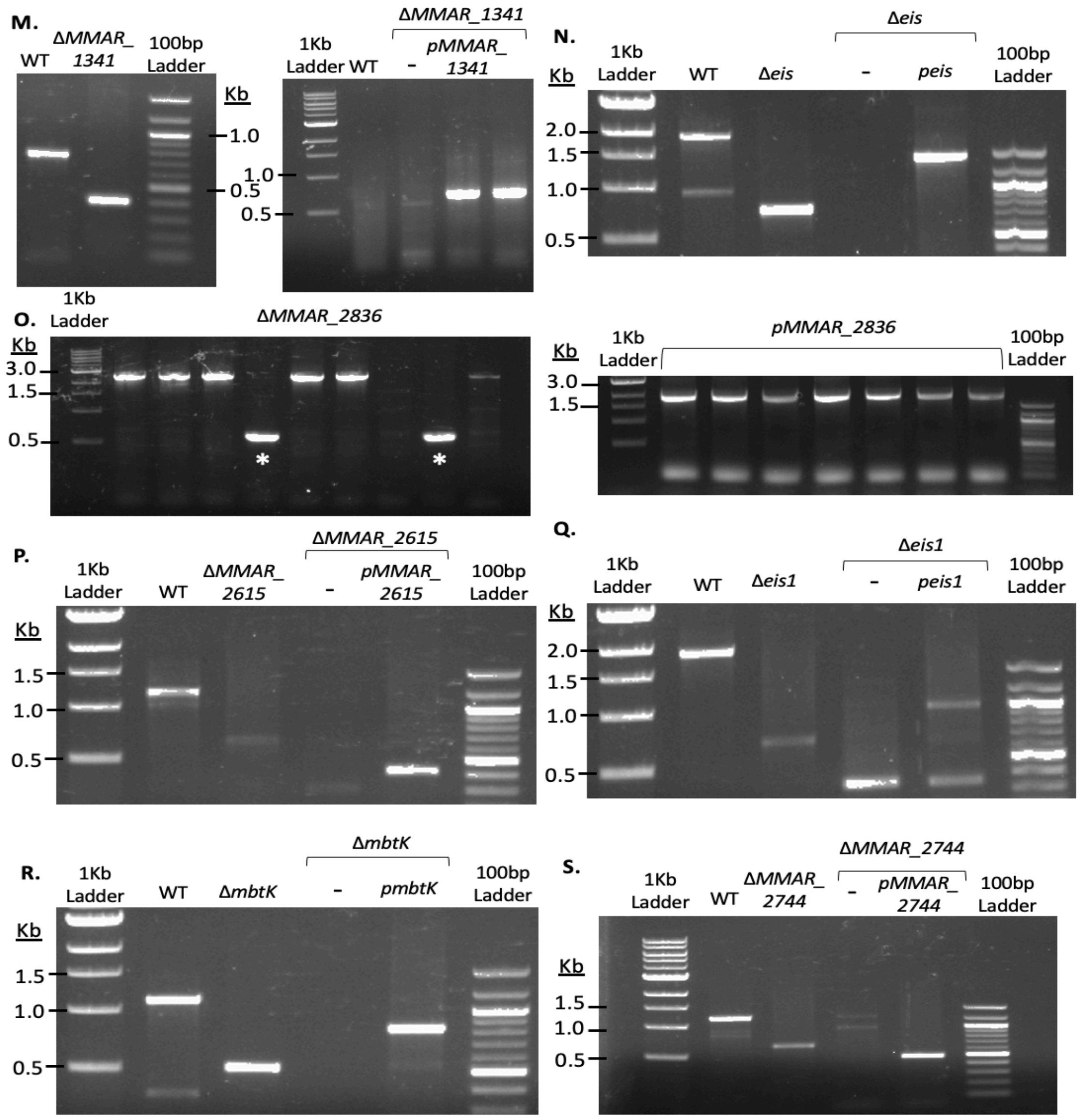


**Figure S1 (Cont.):** ***Mycobacterium marinum*** **Deletion and Complement Strain Confirmations:** PCRs confirming Deletion and complementation of **M.** *MMAR_1341* gene. Product WT: 766bp. Δ*MMAR_1341*: 406 bp. Size plasmid product: 693 bp. Absence in control lanes. **N.** *eis* gene. Product WT: 1,889 bp. Δ*eis*: 695 bp. Size plasmid product: 1,404 bp. Absence in control lane. **O.** *MMAR_2836* gene. Product WT: 2,458 bp. Δ*MMAR_2836*: 619 bp. Size plasmid product: 2,073 bp. **P.** *MMAR_2615* gene. Product WT: 1,182 bp. Δ*MMAR_2615*: 651 bp. Size plasmid product: 450 bp. Absence in control lane. **Q.** *eis1* gene. Product WT: 1,870 bp. Δ*eis1*: 652 bp. Size plasmid product: 1,003 bp. Absence in control lane. **R.** *mbtK* gene. Product WT: 1,099 bp. Δ*mbtK*: 508 bp. Size plasmid product: 816 bp. Absence in control lane. **S.** *MMAR_2744* gene. Product WT: 1,176 bp. Δ*MMAR_2744:* 630 bp. Size plasmid product: 542 bp.

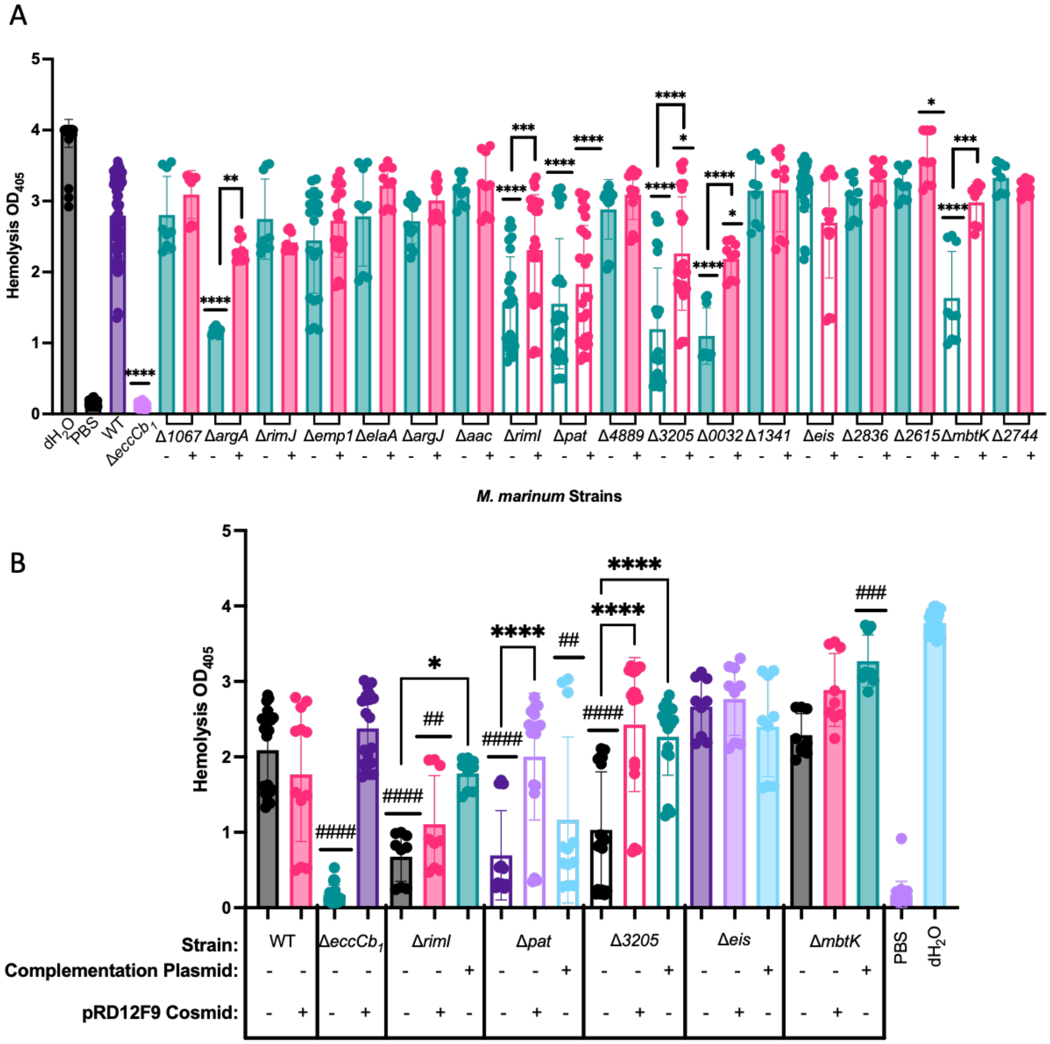


**Figure S2. Hemolytic activity of *M. marinum* strains A.** The majority of the NATs are not required for hemolysis. The putative / known KATs are not filled. -, + refers to absence or presence of complementation plasmid, which gene of interest expressed from the mycobacterial optimal promoter. Significance relative to the WT strain. * $P$=0.0176, ** $P$=0.0068. **B.** The pRD1-2F9 cosmid rescues hemolytic activity of some *M. marinum* KAT deletion strains. Hemolytic activity of *M. marinum* strains lacking genes encoding putative or known KATs. Deletion strains are complemented via expression plasmid behind the mycobacterial optimal promoter, or by pRD1-2F9 cosmid (indicated by – and + symbols, where appropriate). Significance was determined using a one-way ordinary ANOVA ($P$<.0001), followed by a Tukey's multiple comparison test. Significance relative to WT strain reported with # symbols, significance between deletion and complementation strains reported with * symbols. Data is representative of at least 3 biological replicates. #### $P$<0.0001, ### $P$=0.001, ## $P$=0.0048 and 0.0026, **** $P$<0.0001, * $P$=0.0117.

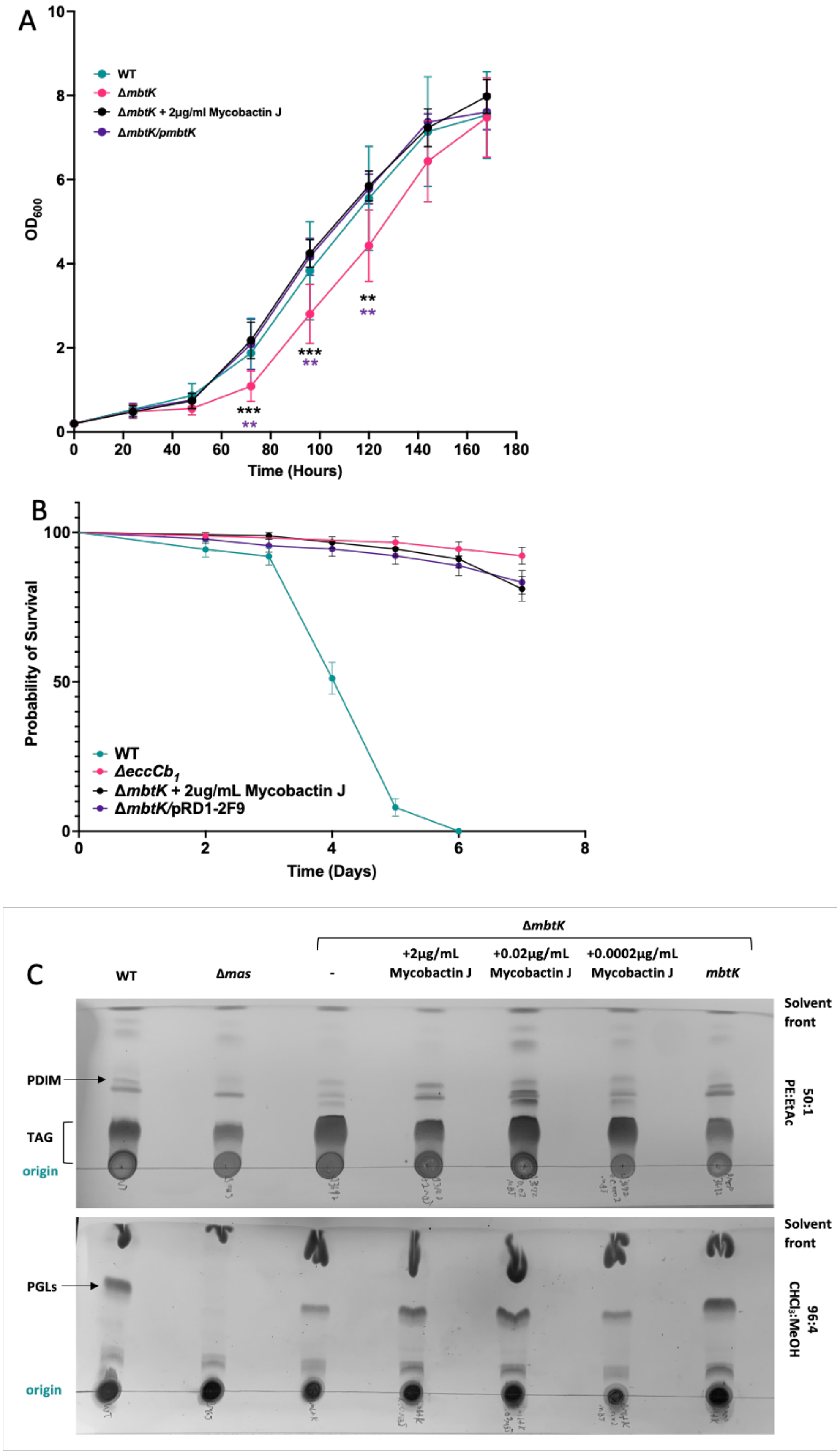
A
WT
ΔmbtK
ΔmbtK + 2µg/ml Mycobactin J
ΔmbtK/pmbtK
OD600
Time (Hours)
B
Probability of Survival
WT
ΔeccCb1
ΔmbtK + 2ug/mL Mycobactin J
ΔmbtK/pRD1-2F9
Time (Days)
C
ΔmbtK
WT
Δmas
-
+2µg/mL Mycobactin J
+0.02µg/mL Mycobactin J
+0.0002µg/mL Mycobactin J
mbtK
Solvent front
PDIM
TAG
origin
50:1 PE:EtAc
PGLs
96:4 CHCl3:MeOH

**Figure S3. Ferric mycobactin J rescues growth and PDIM/PGL production of, but not *G. mellonella* killing by the Δ*mbtK* strain A.** Growth curves of *M. marinum* strains in 7H9 media with 0.1% Tween-80. Strains were grown for 7 days, with growth readings taken every 24 hours by $OD_{600}$. Statistical analysis using a two-way ANOVA. 72h *** $P$=0.0002, ** $P$=0.0043; 96h *** $P$=0.0007, ** $P$=0.0013, 120h ** $P$=0.0037, $P$=0.0052 **B.** *G. mellonella* infection with *M. marinum* strains. Larvae were injected with $1x10^7$ bacteria into the right hindmost proleg. Larvae were infected in 3 groups of 10, for a total of 30 larvae per strain per infection. Data is representative of at least 3 biological replicates, each performed in technical triplicate. Statistical analysis was performed as in (2). Survival curves were compared using a Log-rank Mantel-Cox test, $P$<0.0001 **C.** Thin-layer chromatography of the Δ*mbtK* supplemented with the concentrations of ferric mycobactin J indicated. 0.02 µg/mL mycobactin J was sufficient to restore PDIM and PGL lipid production to WT levels.

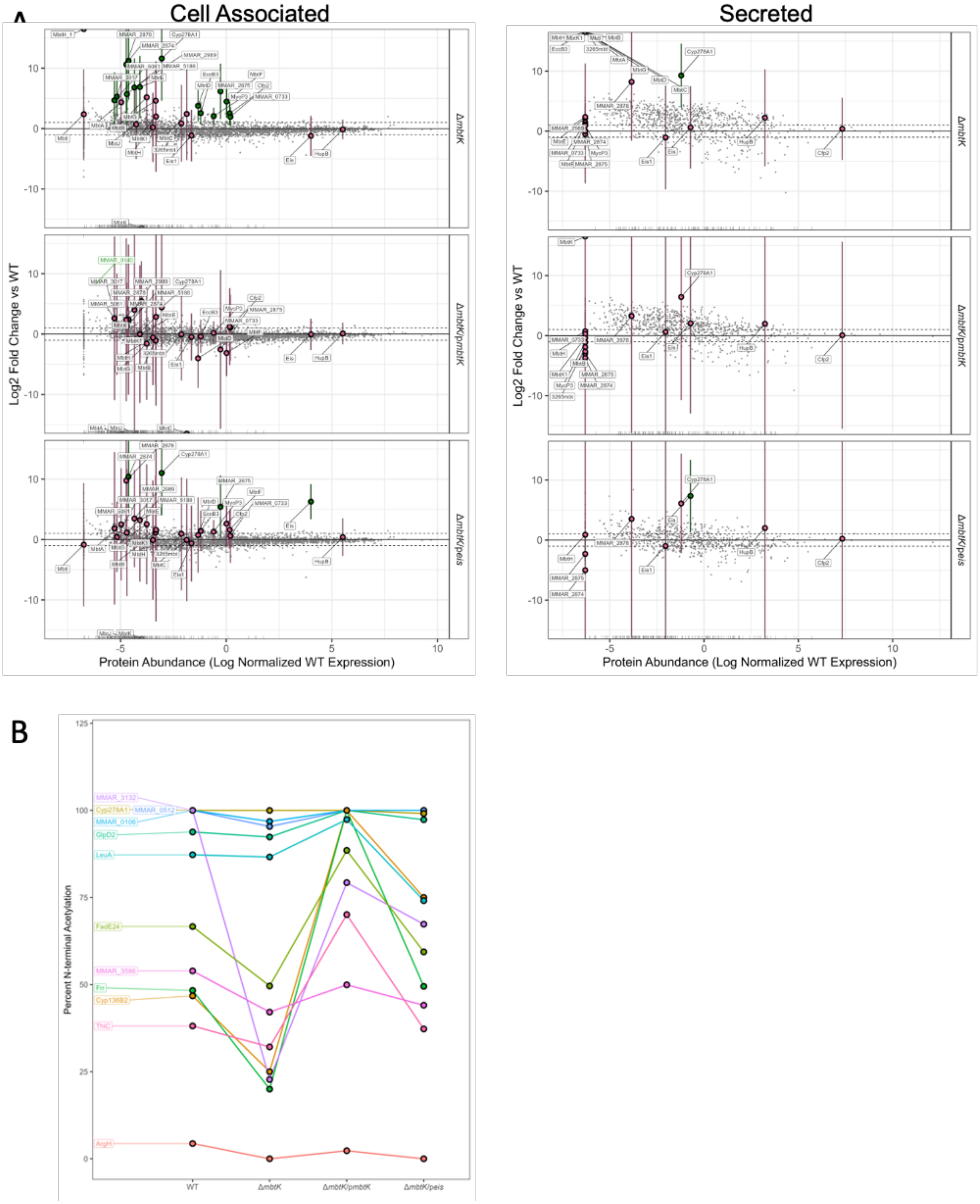
A
Cell Associated
Secreted
Log2 Fold Change vs WT
Protein Abundance (Log Normalized WT Expression)
ΔmbtK
ΔmbtK/pmbtK
ΔmbtK/peis
B
Percent N-terminal Acetylation
MMAR_3132
Cyp278A1
MMAR_0512
MMAR_0106
GlpD2
LeuA
FadE24
MMAR_3586
Frr
Cyp136B2
ThiC
ArgH
WT
ΔmbtK
ΔmbtK/pmbtK
ΔmbtK/peis

**Figure S4: Proteomic measurement of cell associated and secreted protein levels and quantified N-terminal acetylation changes. A.**Proteomic measurement of proteins in the cell associated (left) and secreted (right) in the Δ*mbtK*, Δ*mbtK* complemented with *mbtK (*Δ*mbtK*/p*mbtK*) and the Δ*mbtK* expressing *eis (*Δ*mbtK*/p*eis*) compared to the WT strain. Y-axis is Log2 fold change compared to WT, and x-axis is log normalized WT expression, with more abundant proteins towards the right and less abundant proteins toward the left. Dotted horizontal lines show a log2 fold change of 1 or -1. Large green points are proteins that were significantly changed compared to WT (Benjamini-Hochberg [B-H] corrected *p-value* < 0.05). Large pink points are additional proteins of interest, that did not meet this significance threshold, including proteins that were significant in the WT vs Δ*mbtK* comparison (cell associated), but not in other comparisons. Error bars are 95% confidence intervals, calculated from the B-H corrected *p-values*. Proteins at the bottom of each sub-plot were detected in WT and not in the corresponding mutant strain. Proteins at the top left corner of each sub-plot were detected in the corresponding strain and not WT. Light green points (specifically MMAR_0140) are proteins that were not significantly changed in the Δ*mbtK* vs WT comparison but did meet the significance threshold in Δ*mbtK*/p*mbtK* vs WT. Small grey dots are the remaining quantified proteins in the proteome. **B.** Proteins that showed changing N-terminal acetylation based on quantified N-terminal peptides. Un-acetylated N-termini were acetylated with a deuterated acetyl group (heavy) to compare to the endogenous acetylation (light).Y-axis is percent acetylation calculated by comparing acetylated N-terminal peptide area (light) / (total N-terminal peptide area (light +heavy). Different strains are compared on the x-axis. Proteins were displayed on the plot if the percent acetylation in the Δ*mbtK* strain was less than both the WT and the Δ*mbtK*/p*mbtK* strain, however, the data was too sparse to establish significance.

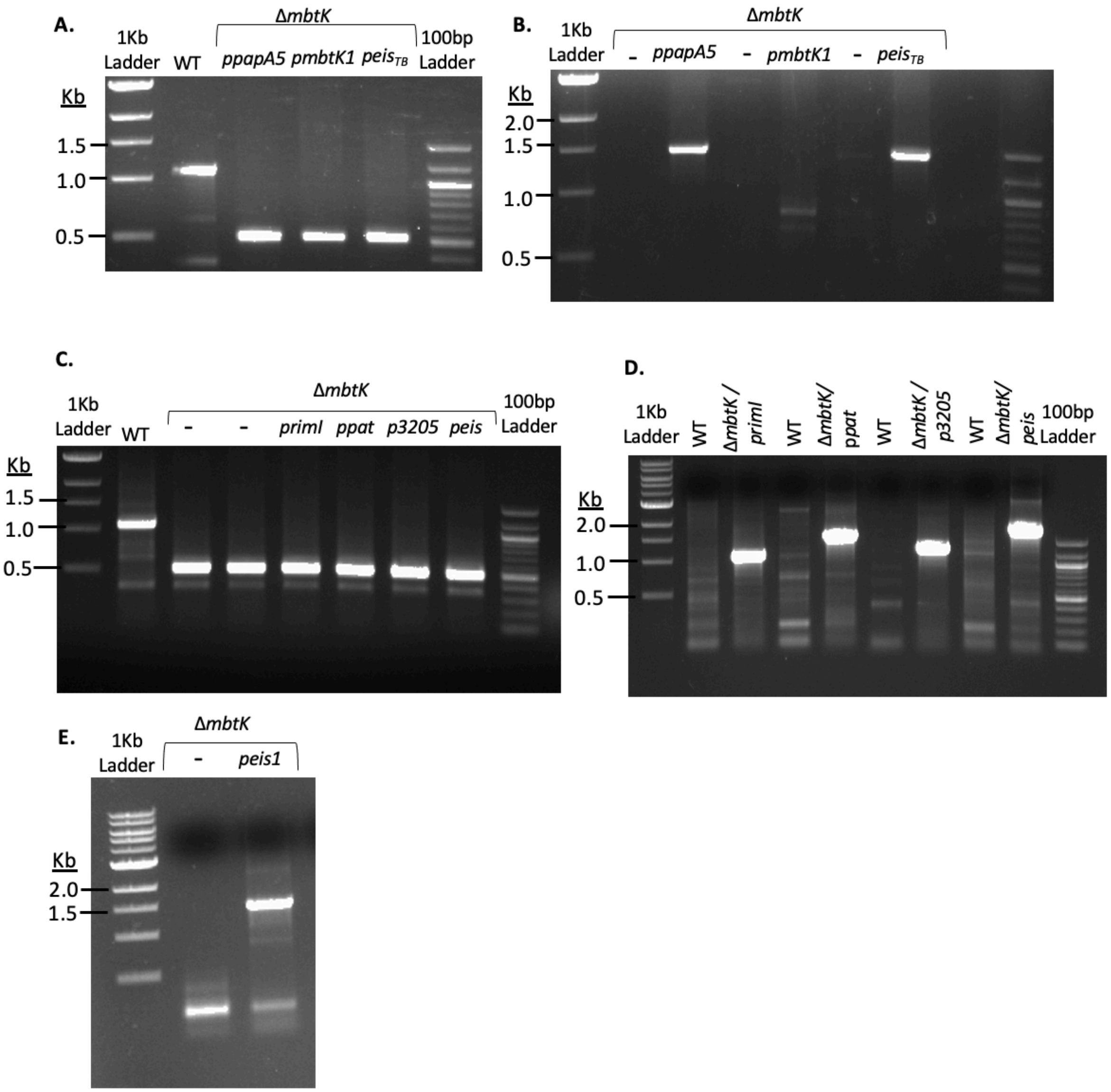


**Figure S5: *Mycobacterium marinum* Cross-Complement Strain Confirmations:** PCRs confirming **A.** deletion of *mbtK* from each cross-complement strain listed. Size of product in WT: 1,099 bp. Δ*mbtK:* 508 bp. **B.** Size of integrating expression plasmid in presence of each cross-complement strain listed: *papA5* 1,450 bp. *mbtK1* 829 bp. $eis_{TB}$ 1,393 bp. **C.** PCRs confirming deletion of *mbtK* from each cross-complement strain listed. Size of product in WT: 1,099 bp. Δ*mbtK:* 508 bp. **D.** Size of integrating expression plasmid in each cross-complement strain: *primI* 1,088 bp. *ppat* 1,562 bp. *MMAR_3205* 1,271 bp. *peis* 1,829 bp. **E.** Size of *eis1* integrating expression plasmid in Δ*mbtK/peis1* strain: 1,423 bp.

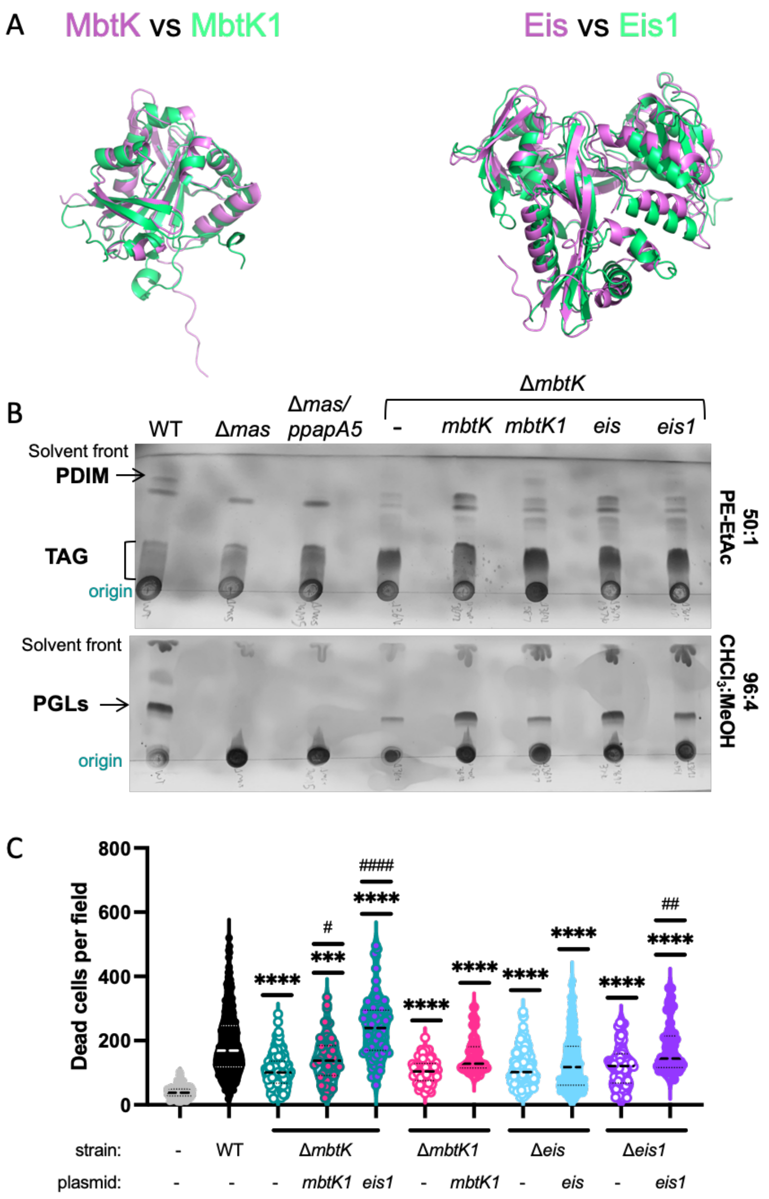


**Figure S6: Cross-Complementation of Δ*mbtK* with *mbtK1* and *eis1*. A.** Alignments of MbtK with MbtK1 and Eis with Eis1. Protein structures obtained from AlphaFold (3), and proteins visualized on PyMOL (4). **B.** Thin-layer chromatography of the Δ*mbtK* strain expressing the *mbtK, mbtK1, eis,* and *eis1* genes. This TLC is representative of at least three biological replicates. **C.** RAW 264.7 macrophage infections with *M. marinum* strains bearing deletions in the *eis, eis1, mbtK* and *mbtK1* genes, and corresponding complementation and cross complementation strains. The experiment includes three biological replicates, each performed in triplicate, with 15 images counted per well. Outliers were removed using ROUT analysis, with a Q value of 0.5%, using Prism 10. Statistical significance was determined using a one-way ANOVA ($P<0.0001$) followed by a Tukey's multiple comparison test. Significance vs the WT strain is indicated using *. **** $P<0.0001$, *** $P=0.0008$. Significance vs the Δ strain is indicated using #. #### $P<0.0001$, ## $P=0.0023$, # $P=0.0437$.

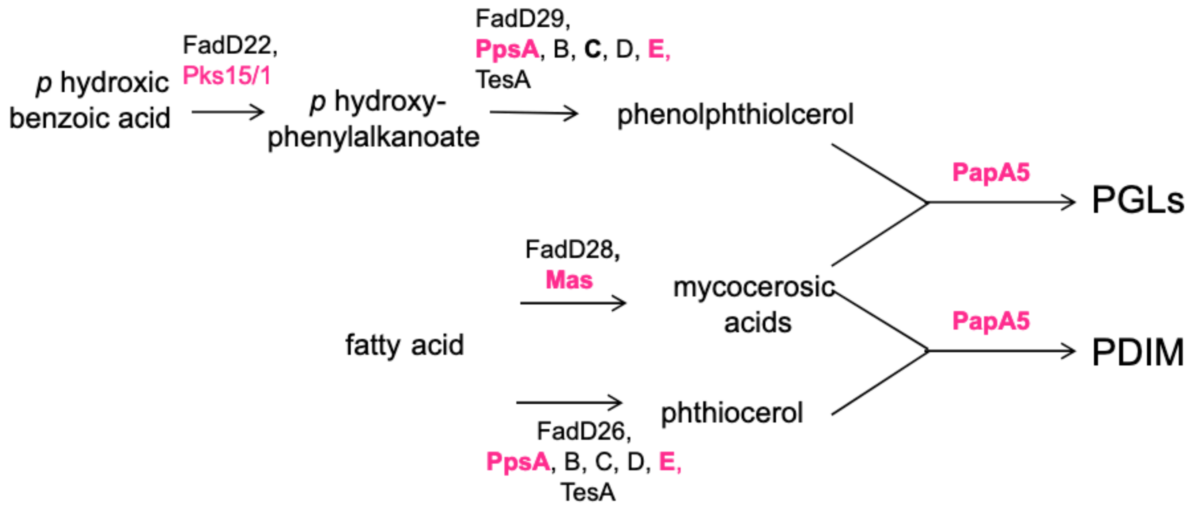


**Figure S7: PDIM and PGL Biosynthetic Pathway** Graphical representation of proteins facilitating PDIM and PGL biosynthesis. Proteins highlighted in pink have been previously identified as post-translationally acetylated.

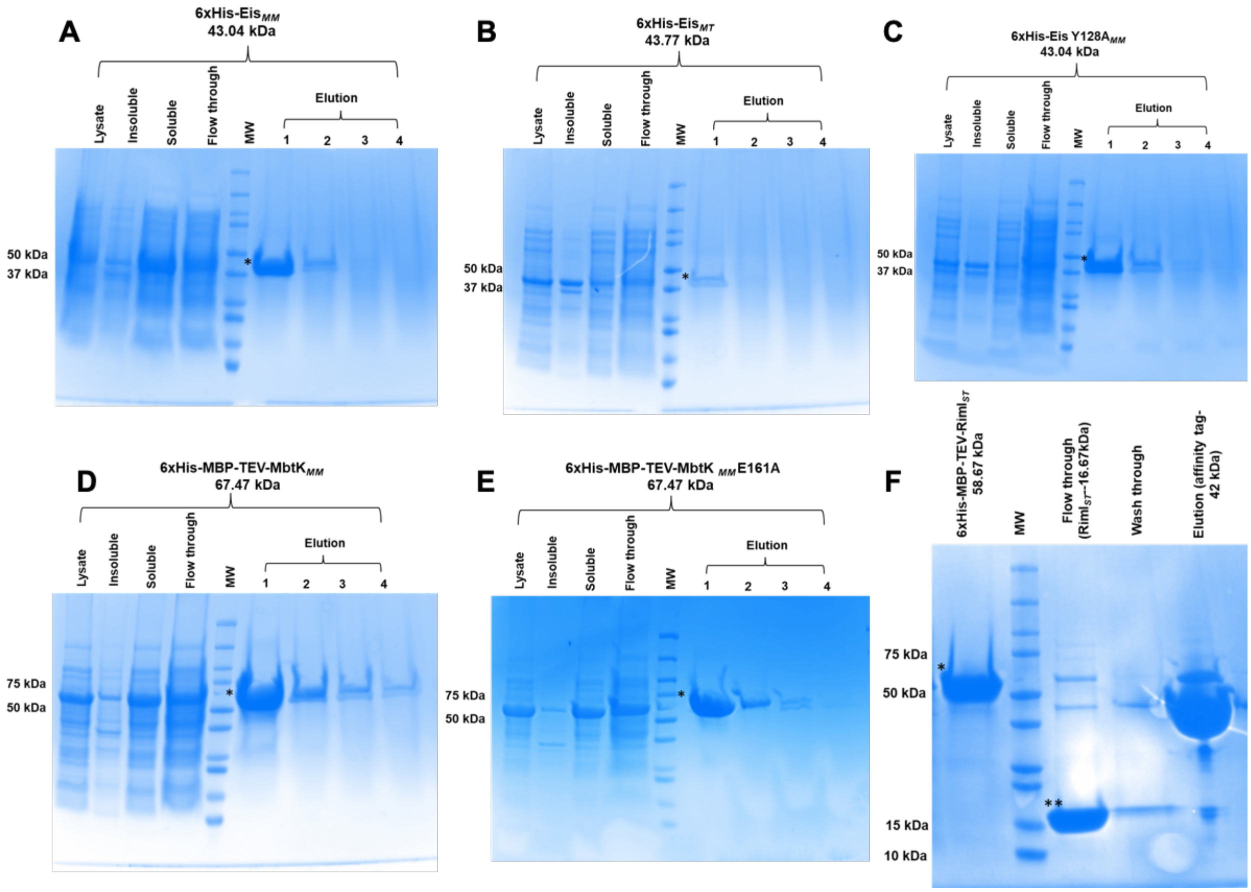


**Figure S8: Heterologous expression of NAT proteins.** Coomassie-stained gels representing purification of 6×His-Eis$_{MM}$ protein (A) and 6×His-Eis Y128A$_{MM}$ protein (B) from *M. marinum* and 6×His-Eis$_{MT}$ protein (C) from *M. tuberculosis*. (D) Purification of 6×His-MbtK$_{MM}$ protein with MBP tag from *M. marinum.* (E) Coomassie-stained gel represents purified 6×His-MBP-TEV-RimI$_{ST}$ protein from *Salmonella typhimurium* (marked with *), further purified by affinity tag removal using 6×His-TEV protease.(F) Purification of 6×His-MbtK$_{MM}$E161A protein with MBP tag from *M. marinum.* Final products were obtained by a secondary incubation with Ni-NTA resin to remove protease, tag and uncut proteins (marked with **), present in the flow through. Final concentrations and buffer conditions are listed in the methods section.

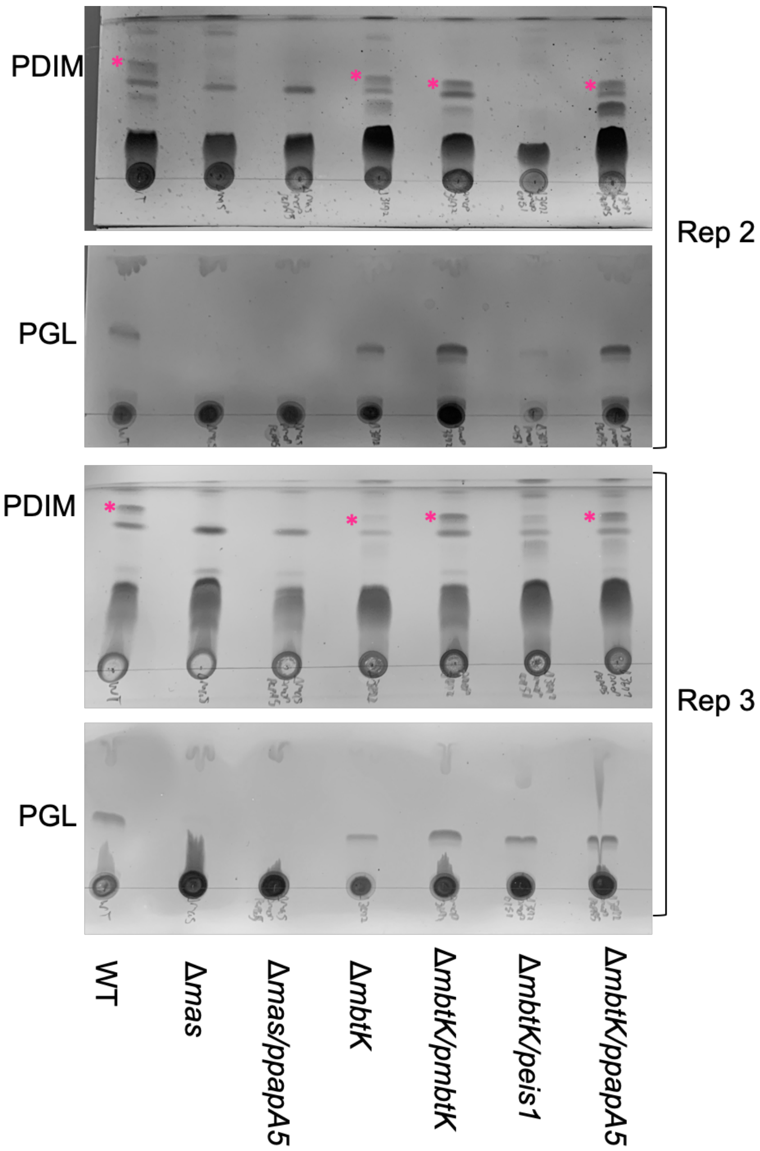


**Figure S9: PapA5 restores PDIM/PGL to the Δ*mbtK* strain.** Thin-layer chromatography of the *M. marinum* strains. These TLCs represent additional replicates of the data found in Figure 3F. Pink asterisk signifies the PDIM band. Performed exactly as Fig. S3F.

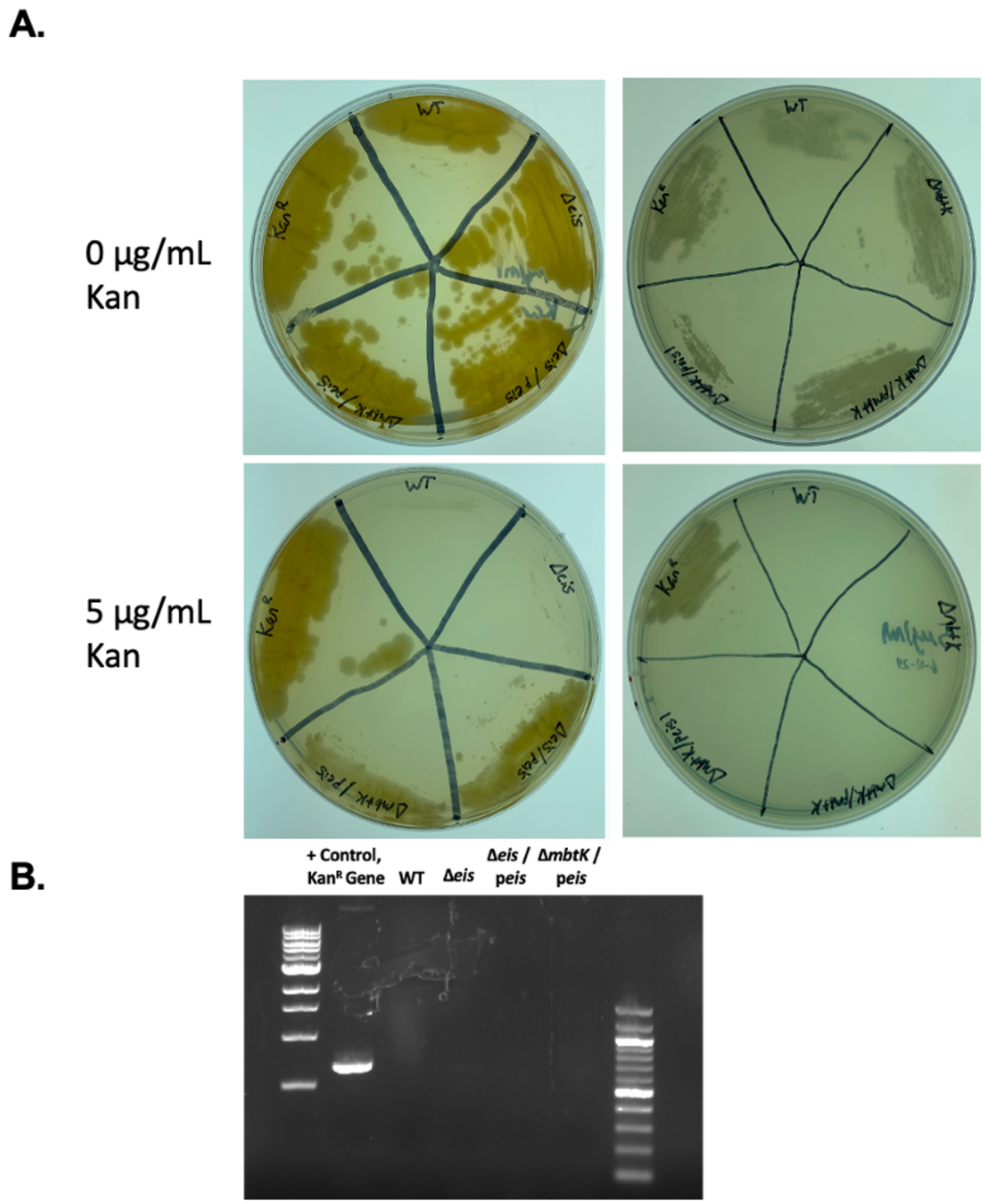


**Figure S10: Kanamycin resistance capacity of strains constitutively expressing *eis*. A.** Growth of  . *marinum* strains on 7H11 agar with kan concentrations up to 5 μg/mL. **B.** PCR of the kanamycin resistance gene. The kanamycin resistance gene is not present in the *eis* deletion strain, or in strains expressing *eis* via integrating plasmid. Expected sizes: 686 bp if gene is present, absence of band if gene is not present within a strain, respectively.

**Table S1. Bacterial strains used in this study**

| Name | Genotype | Reference |
|---|---|---|
| DH5α | *E. coli* F⁻ φ80*lac*ZΔM15 Δ(*lac*ZYA-*arg*F)U169 *rec*A1 *end*A1 *hsd*R17($r_K^-$, $m_K^+$) *pho*A *sup*E44 $\lambda^-$ *thi*-1 *gyr*A96 *rel*A1 | NEB |
| *M. marinum* M strain | Wild type, parental strain | ATCC BAA-535 |
| Δ$eccCb_1$ | M strain with an unmarked in-frame deletion of the $eccCb_1$ gene | (5) |
| Δ*mas* | M strain with an unmarked in-frame deletion of the *mas* gene | (6) |
| Δ*MMAR_1067* | M strain with an unmarked in-frame deletion of the *MMAR_1067* gene | (6) |
| Δ*argA* | M strain with an unmarked in-frame deletion of the *argA* gene | (6) |
| Δ*rimJ* | M strain with an unmarked in-frame deletion of the *rimJ* gene | (6) |
| Δ*emp1* | M strain with an unmarked in-frame deletion of the *emp1* gene | (6) |
| Δ*elaA* | M strain with an unmarked in-frame deletion of the *elaA* gene | (6) |
| Δ*argJ* | M strain with an unmarked in-frame deletion of the *argJ* gene | This study |
| Δ*aac* | M strain with an unmarked in-frame deletion of the *aac* gene | This study |
| Δ*rimI* | M strain with an unmarked in-frame deletion of the *rimI* gene | This study |
| Δ*pat* | M strain with an unmarked in-frame deletion of the *pat* gene | This study |
| Δ*MMAR_4889* | M strain with an unmarked in-frame deletion of the *MMAR_4889* gene | This study |
| Δ*MMAR_3205* | M strain with an unmarked in-frame deletion of the *MMAR_3205* gene | This study |
| Δ*MMAR_0332* | M strain with an unmarked in-frame deletion of the *MMAR_0332* gene | This study |
| Δ*MMAR_1341* | M strain with an unmarked in-frame deletion of the *MMAR_1341* gene | This study |
| Δ*eis* | M strain with an unmarked in-frame deletion of the *eis* gene | This study |
| Δ*MMAR_2836* | M strain with an unmarked in-frame deletion of the *MMAR_2836* gene | This study |
| Δ*MMAR_2615* | M strain with an unmarked in-frame deletion of the *MMAR_2615* gene | This study |
| Δ*eis1* | M strain with an unmarked in-frame deletion of the *eis1* gene | This study |
| Δ*mbtK* | M strain with an unmarked in-frame deletion of the *mbtK* gene | This study |
| Δ*MMAR_2744* | M strain with an unmarked in-frame deletion of the *MMAR_2744* gene | This study |

**Table S2 Plasmids used in this study**

| Name | Genotype | Reference |
|---|---|---|
| p2NIL | Parental suicide vector for allelic exchange, *kan*$^R$, *amp*$^R$ . | Addgene plasmid #20188. (7) |
| pGOAL19 | Marker cassette for allellic exchange, *amp*$^R$, *hyg*$^R$, *lacZ, sacB.* | Addgene plasmid #20190 (7) |
| pET15b | Expression vector with T7 promoter and N-terminal 6xHis tag, *amp*$^R$. | Novagen |
| PET28-MBP-TEV | Expression vector with T7 promoter for bacterial expression of 6xHis-MBP-TEV N-terminal fusions, *kan*$^R$. | Addgene plasmid # 69929 (8) |
| p2NILΔ*argJGOAL* | Allelic exchange plasmid to generate the Δ*argJ* strain. p2NIL backbone with the GOAL19 marker cassette. Contains *M. marinum argJ* flanking regions, *kan*$^R$, *hyg*$^R$, *lacZ, sacB*. | This study |
| p2NILΔ*aacGOAL* | Allelic exchange plasmid to generate the Δ*aac* strain. p2NIL backbone with the GOAL19 marker cassette. Contains *M. marinum aac* flanking regions, *kan*$^R$, *hyg*$^R$, *lacZ, sacB*. | This study |
| p2NILΔ*rimIGOAL* | Allelic exchange plasmid to generate the Δ*rimI* strain. p2NIL backbone with the GOAL19 marker cassette. Contains *M. marinum rimI* flanking regions, *kan*$^R$, *hyg*$^R$, *lacZ, sacB*. | This study |
| *p2NILΔpatGOAL* | Allelic exchange plasmid to generate the Δ*pat* strain. p2NIL backbone with the GOAL19 marker cassette. Contains *M. marinum pat* flanking regions, *kan*$^R$, *hyg*$^R$, *lacZ, sacB*. | This study |
| p2NILΔ*4889GOAL* | Allelic exchange plasmid to generate the Δ*MMAR_4889* strain. p2NIL backbone with the GOAL19 marker cassette. Contains *M. marinum 4889* flanking regions, *kan*$^R$, *hyg*$^R$, *lacZ, sacB*. | This study |
| p2NILΔ*3205GOAL* | Allelic exchange plasmid to generate the Δ*MMAR_3205* strain. p2NIL backbone with the GOAL19 marker cassette. Contains *M. marinum 3205* flanking regions. *kan*$^R$, *hyg*$^R$, *lacZ, sacB*. | This study |
| p2NILΔ*0332GOAL* | Allelic exchange plasmid to generate the Δ*MMAR_0332* strain. p2NIL backbone with the GOAL19 marker cassette. Contains *M. marinum 0332* flanking regions, *kan*$^R$, *hyg*$^R$, *lacZ, sacB*. | This study |
| p2NILΔ*1341GOAL* | Allelic exchange plasmid to generate the Δ*MMAR_1341* strain. p2NIL backbone with the GOAL19 marker cassette. Contains *M. marinum 1341* flanking regions, *kan*$^R$, *hyg*$^R$, *lacZ, sacB*. | This study |
| p2NILΔ*eisGOAL* | Allelic exchange plasmid to generate the Δ*eis* strain. p2NIL backbone with the GOAL19 | This study |

| | | |
|---|---|---|
| | marker cassette. Contains *M. marinum eis* flanking regions. *kan*$^R$, *hyg*$^R$, *lacZ, sacB*. | |
| p2NILΔ*2836GOAL* | Allelic exchange plasmid to generate the Δ*MMAR_2836* strain. p2NIL backbone with the GOAL19 marker cassette. Contains *M. marinum 2836* flanking regions, *kan*$^R$, *hyg*$^R$, *lacZ, sacB*. | This study |
| p2NILΔ*2615GOAL* | Allelic exchange plasmid to generate the Δ*MMAR_2615* strain. p2NIL backbone with the GOAL19 marker cassette. Contains *M. marinum 2615* flanking regions, *kan*$^R$, *hyg*$^R$, *lacZ, sacB*. | This study |
| p2NILΔ*eis1GOAL* | Allelic exchange plasmid to generate the Δ*eis1* strain. p2NIL backbone with the GOAL19 marker cassette. Contains *M. marinum eis1* flanking regions, *kan*$^R$, *hyg*$^R$, *lacZ, sacB*. | This study |
| p2NILΔ*mbtKGOAL* | Allelic exchange plasmid to generate the Δ*mbtK* strain. p2NIL backbone with the GOAL19 marker cassette. Contains *M. marinum mbtK* flanking regions, *kan*$^R$, *hyg*$^R$, *lacZ, sacB*. | This study |
| p2NILΔ*2744GOAL* | Allelic exchange plasmid to generate the Δ*MMAR_2744* strain. p2NIL backbone with the GOAL19 marker cassette. Contains *M. marinum 2744* flanking regions, *kan*$^R$, *hyg*$^R$, *lacZ, sacB*. | This study |
| pRD1-2F9 | Plasmid containing an extended RD1 region from *M. tuberculosis*. *hyg*$^R$. | (9) |
| p*MMAR_1067* | *MMAR_1067* expressed behind the mycobacterial optimal promoter ($p_{MOP}$), integrated at *attB, hyg*$^R$. | This study |
| p*argA* | *argA* expressed behind the mycobacterial optimal promoter integrated at *attB, hyg*$^R$. | (6) |
| p*rimJ* | *rimJ* expressed behind the mycobacterial optimal promoter integrated at *attB, hyg*$^R$. | This study |
| p*emp1* | *MMAR_1839* expressed behind the mycobacterial optimal promoter integrated at *attB, hyg*$^R$. | This study |
| p*elaA* | *elaA* expressed behind the mycobacterial optimal promoter integrated at *attB, hyg*$^R$ . | This study |
| p*argJ* | *argJ* expressed behind the mycobacterial optimal promoter integrated at *attB, hyg*$^R$. | This study |
| p*aac* | *aac* expressed behind the mycobacterial optimal promoter integrated at *attB, hyg*$^R$. | This study |
| p*rimI* | *rimI* expressed behind the mycobacterial optimal promoter integrated at *attB, hyg*$^R$. | This study |
| *ppat* | *pat* expressed behind the mycobacterial optimal promoter integrated at *attB, hyg*$^R$. | This study |
| p*MMAR_4889* | *MMAR_4889* expressed behind the mycobacterial optimal promoter ($p_{MOP}$), integrated at *attB, hyg*$^R$. | This study |
| p*MMAR_3205* | *MMAR_3205* expressed behind the mycobacterial optimal promoter integrated at *attB, hyg*$^R$. | This study |

| | | |
|---|---|---|
| p*MMAR_0332* | *MMAR_0332* expressed behind the mycobacterial optimal promoter integrated at *attB, hyg*$^R$. | This study |
| p*MMAR_1341* | *MMAR_1341* expressed behind the mycobacterial optimal promoter integrated at *attB, hyg*$^R$. | This study |
| p*eis* | *eis* expressed behind the mycobacterial optimal promoter integrated at *attB, hyg*$^R$. | This study |
| p*eisY128A* | *eis* expressed behind the mycobacterial optimal promoter integrated at *attB, hyg*$^R$. Y residue at position 128 changed to an A. | This study |
| p*ERDMAN_2659* | *ERDMAN_2659* expressed behind the mycobacterial optimal promoter integrated at *attB, hyg*$^R$. | This study |
| p*MMAR_2836* | *MMAR_2836* expressed behind the mycobacterial optimal promoter integrated at *attB, hyg*$^R$. | This study |
| p*MMAR_2615* | *MMAR_2615* expressed behind the mycobacterial optimal promoter integrated at *attB, hyg*$^R$. | This study |
| p*eis1* | *eis1* expressed behind the mycobacterial optimal promoter integrated at *attB, hyg*$^R$. | This study |
| p*mbtK* | *mbtK* expressed behind the mycobacterial optimal promoter integrated at *attB, hyg*$^R$. | This study |
| p*mbtKE161A* | *mbtK* expressed behind the mycobacterial optimal promoter integrated at *attB, hyg*$^R$. E residue at position 161 changed to an A. | This study |
| p*MMAR_2744* | *MMAR_2744* expressed behind the mycobacterial optimal promoter integrated at *attB, hyg*$^R$. | This study |
| p*papa5* | *papa5* expressed behind the mycobacterial optimal promoter integrated at *attB, hyg*$^R$. | This study |
| p*mbtK1* | *mbtK1* expressed behind the mycobacterial optimal promoter integrated at *attB, hyg*$^R$. | This study |
| pET15b_$eis_{MT}$ | *ERDMAN_2659 M. tuberculosis* Erdman strain with N terminal 6x His tag, a*mp*$^R$. | This study |
| pET15b_*eisY128A* | *eisY128A M. marinum* with N terminal 6x His tag, a*mp*$^R$. | This study |
| pET15b_*eis* | *eis M. marinum* with N terminal 6x His tag, a*mp*$^R$. | This study |
| pET28_*mbtK* | *mbtK M. marinum* with N terminal 6x His-MBP-TEV, k*an*$^R$. | This study |
| pET28_*mbtKE161A* | *mbtKE161A M. marinum* with N terminal 6x His-MBP-TEV, k*an*$^R$. | This study |
| pET28_$RimI_{ST}$ | *rimI S. typhimurium with* with N terminal 6x His-MBP-TEV, k*an*$^R$. | This study |

**Table S3 Primers used in this study**

| Primer name | Sequence 5'-3' | Application |
|---|---|---|
| OVP93 | TGACAGATTACTTAAAGAACCTTCTAG | Amplification of pET28b-MBP-TEV vector |
| OVP94 | GTAGCAGCCTGTACTGAGG | |
| OMF512 | ACTAGTAACTAGCATAACCCCTTGGGGC | Amplification of pET15b vector |
| OMF513 | CATATGATGATGATGATGATGGCTGC | |
| OVP43 | TCATCATCATCATATGACTGCAGTACTGCCCGCATC | Amplification of *eis* and *eisY128A* |
| OVP44 | TATGCTAGTTACTAGTCTAGAACTCGAAAGCCGTCTCGG | |
| OVP99 | TCATCATCATCATATGACTGTGACCCTGTGTAG | Amplification of $eis_{MT}$ |
| OVP100 | TATGCTAGTTACTAGTTCAGAACTCGAACGCGGT | |
| OVP95 | AGTACAGGCTGCTACATGGCGGGCCATCCGGG | Amplification of *mbtK* and *mbtKE161A* |
| OVP96 | CTTTAAGTAATCTGTCATCATCGGCGTTGCCGGTCAC | |
| OMF088 | GGCTTAAGTATAAGGAGGAAAACATATGAACACGATTTCTATCCTCAGCAC | Amplification of pET23a-RimIST vector |
| OMF089 | CTCCGTTTAGAGAGGGGTTATACTAGTTACATGCTTATCGGTAACGCCATG | |
| OMF310 | ATCTTTATTTTCAAGGAAACACGATTTCTATCCTCAGCAC | Amplification of *rimI* from *Salmonella typhimurium* |
| OMF311 | TTAGCAGCCGGATCTCACATGCTTATCGGTAACGCC | |
| OMF033 | TCCTTGAAAATAAAGATTTTCGGATCCGGATTGGAAGTACAGG | Amplification of pET28bMBP vector (Gift from Dr. Shaun Lee) |
| OMF034 | TGAGATCCGGCTGCTAACAAAG | |

| | | |
|---|---|---|
| OMF283 | CGTGGTGTCACGCTCGTGGGAGGACTGTTAGGTCTGATTGGCACC | PCR primers amplifying flanking regions of *argJ* to facilitate deletion. This study. |
| OMF284 | GTCCGCTTAAGAGTAATGCCTTGACTACGCAACAGCCGC | |
| OMF285 | ATTACTCTTAAGCGGACCACCGATCTGTCCCACG | |
| OMF286 | ACGCAGTCAGGCACCGTTGCGCGTAGAAGGCTCCGGTGC | |
| OMF291 | CGTGGTGTCACGCTCGTGATATCTCAACCGAGCTTGCGGCGG | PCR primers amplifying flanking regions of *aac* to facilitate deletion. This study. |
| OMF292 | CATCAACTTAAGCAGCCGAGCCGTGTGAACCTCG | |
| OMF293 | GGCTGCTTAAGTTGATGTGCGACTGGCGCGC | |
| OMF294 | ACGCAGTCAGGCACCGTAGCAATTGCTGCCGCATTGCCC | |
| OMF357 | CGTGGTGTCACGCTCGTCCGTTGGGCGAGTTCCTGAAAAACCC | PCR primers amplifying flanking regions of *rimI* to facilitate deletion. This study. |
| OMF358 | CGCTGACCTTAAGGATGGTGACGGGCCCGAGATCG | |
| OMF359 | ACCATCCTTAAGGTCAGCGGTGCCGACGCATACACG | |
| OMF360 | ACGCAGTCAGGCACCGTGCTTCTCGCCGTCCTCGTCCCAGC | |
| OMF373 | CGTGGTGTCACGCTCGTCCTGGAAAAGATGTACCAGGCCC | PCR primers amplifying flanking regions of *pat* to facilitate deletion. This study. |
| OMF374 | TGGCGCTTAAGGGTCACTTCGGCCATCCCGTCC | |
| OMF375 | GTGACCCTTAAGCGCCAGGTGATCGAGGCGGTG | |
| OMF376 | ACGCAGTCAGGCACCGTCGCATGGGGATCTGCTGGTGATGG | |
| OMF393 | CGTGGTGTCACGCTCGTAACGCGATACCGCGACCGGTGG | PCR primers amplifying flanking regions of |

| | | |
|---|---|---|
| OMF394 | CAGCTCCTTAAGCGGCCAGTGACGAGACATCGCCC | *MMAR_4889* to facilitate deletion. This study. |
| OMF395 | TGGCCGCTTAAGGAGCTGTTCGGCGTTTCGCGC | |
| OMF396 | ACGCAGTCAGGCACCGTAACTGAGCGAGATAGACCGCTGGCCG | |
| OMF403 | CGTGGTGTCACGCTCGTTTCGGGCAGAGACCACGGTGC | PCR primers amplifying flanking regions of *MMAR_3205* to facilitate deletion. This study. |
| OMF404 | GCGCGCTTAAGCAGATCAATGACGAAGATCGTCAACGCG | |
| OMF405 | GATCTGCTTAAGCGCGCGCTTCCCCTGTAGGCC | |
| OMF406 | ACGCAGTCAGGCACCGTGGATGATCGCCGAAGACGACGATCG | |
| OMF411 | CGTGGTGTCACGCTCGTCTGTGTGGCAGAAAGCGTCGTCCG | PCR primers amplifying flanking regions of *MMAR_0332* to facilitate deletion. This study. |
| OMF412 | CACATCTTAAGCTCACGGATATCGGCCTTCTGCGC | |
| OMF413 | CGTGAGCTTAAGATGTGGCGTGACCCGCGGTAAGC | |
| OMF414 | ACGCAGTCAGGCACCGTCTGGGCGTCGACTCGCTGATGTCG | |
| OMF419 | CGTGGTGTCACGCTCGTACCGACATCGAGCTGACCGAAACCG | PCR primers amplifying flanking regions of *MMAR_1341* to facilitate deletion. This study. |
| OMF420 | TGCCCCTTAAGAATTTCAGCGGTGTCTTCCGGCTTGG | |
| OMF421 | GAAATTCTTAAGGGGCAGCCGCAGAACGAATGG | |
| OMF422 | ACGCAGTCAGGCACCGTGAGCGCGTGGGCTATCTCACTTTGG | |
| OBJ121 | CGTGGTGTCACGCTCGTGCGGCAAACTTGTACAGCAC | PCR primers amplifying flanking regions of *eis* to facilitate deletion. This study. |
| OBJ122 | CTAGAACTTAAGAGTCACGGGCTCCGACTCTG | |

| | | |
|---|---|---|
| OBJ123 | GTGACTCTTAAGTTCTAGGTTCCGCCGCGCTG | |
| OBJ124 | GCAGTCAGGCACCGTACCTGGTCAGGCTCTAACTC | |
| OBJ129 | CGTGGTGTCACGCTCGTATCCCACAGGGTGCATCATC | PCR primers amplifying flanking regions of *MMAR_2836* to facilitate deletion. This study. |
| OBJ130 | CGAGCCCTTAAGGACCATTCAGCGATTGTTGC | |
| OBJ131 | ATGGTCCTTAAGGGCTCGTGGTGAGTGCTAGTG | |
| OBJ132 | GCAGTCAGGCACCGTCACGCTGAACGTCTCGATAC | |
| OBJ137 | CGTGGTGTCACGCTCGTTCAGATAGCCGTCCGGATCG | PCR primers amplifying flanking regions of *MMAR_2615* to facilitate deletion. This study. |
| OBJ138 | GATTCACTTAAGCACGCTTGCTCCCATGGTTTG | |
| OBJ139 | AGCGTGCTTAAGTGAATCCTGACCTCGCCGTG | |
| OBJ140 | GCAGTCAGGCACCGTCCGTAGGCAAGCCCAAGAAC | |
| OBJ145 | CGTGGTGTCACGCTCGTGCGTGTACAGCTGCAGGATG | PCR primers amplifying flanking regions of *eis1* to facilitate deletion. This study. |
| OBJ146 | GCCCTACTTAAGGCTCACCCGGCCATCATGTCG | |
| OBJ147 | GTGAGCCTTAAGTAGGGCGTACTGGCGTTCTACC | |
| OBJ148 | GCAGTCAGGCACCGTGGGTGCGCTCAAAGGTCTTC | |
| OBJ153 | CGTGGTGTCACGCTCGTTCAACGAGAGCGGAAGATGC | PCR primers amplifying flanking regions of *mbtK* to facilitate deletion. This study. |
| OBJ154 | TCGGCGCTTAAGCGCCATCAGCGCCTGGG | |
| OBJ155 | ATGGCGCTTAAGCGCCGATGAGCCCAACCG | |
| OBJ156 | GCAGTCAGGCACCGTGGGCATTGAGTCTCGCGAAC | |
| OBJ161 | CGTGGTGTCACGCTCGTATGCCAGCTTGTCGCCAGTG | PCR primers amplifying flanking regions of |

| | | |
|---|---|---|
| OBJ162 | CGGCTACTTAAGGGCCATTCAATCGGCGTGC | *MMAR_2744* to facilitate deletion. This study. |
| OBJ163 | ATGGCCCTTAAGTAGCCGCCCGGCGCGGAATC | |
| OBJ164 | GCAGTCAGGCACCGTAGGCCGAGCGCACGATGATTACG | |
| OGC3 | CCGGTCTGGAATCGTCAATC | Verification of Δ*MMAR_1067* strain. (6) |
| OGC4 | CGCTGGGTCAGATTGGGATG | |
| OGC5 | ATTCGGTGGCCATGCCGTAG | Verification of Δ*argA* strain. (6) |
| OGC6 | TGGTGGTGTCCAGTCAGTTC | |
| OGC7 | TGGGCAAACGCCAACCGATG | Verification of Δ*rimJ* strain. (6) |
| OGC8 | GCATGGGTACCAGCACAAAC | |
| OGC9 | CGTCAACGGTCCTGGTGAAG | Verification of Δ*emp1* strain. (6) |
| OGC10 | AGAACAGTGACGCCAAGGAG | |
| OGC11 | TCAGCGAGTGCAGGTGATCC | Verification of Δ*elaA* strain. (6) |
| OGC12 | TGTTCGCCATCGCGCAAGAC | |
| OGC13 | ATCTATTTGATGCCGGAGGG | Verification of Δ*argJ* strain. This study. |
| OGC14 | AGGACTTCCGGTGTGGTGAC | |
| OGC15 | TGACGCACCGCAATCCAGAC | Verification of Δ*aac* strain. This study. |
| OGC16 | TCCCAGTGCTCGATCCGATG | |
| OGC17 | TCGATGCCATCGGTGGACTC | Verification of Δ*rimI* strain. This study. |
| OGC18 | CCGAACCGCACATGCTCATC | |

| | | |
|---|---|---|
| OMF377 | TGCCCATTCTGATGCCGAGTCCTGC | Verification of Δ*pat* strain. This study. |
| OMF378 | CAGGATGGCCAGCAGACCGATCC | |
| OMF397 | TGCGACTGTGTCAGTCTGGTGGC | Verification of Δ*MMAR_4889* strain. This study. |
| OMF398 | GACGGTGAAGTGTCACGGCTGG | |
| OMF407 | CGATGCAACCCATCACAATTGCCCCG | Verification of Δ*MMAR_3205* strain. This study. |
| OMF408 | CATTGCGATTGCCAGCAACCGCC | |
| OMF415 | CGATGATGGAGCCACCCCATCGTAAGG | Verification of Δ*MMAR_0332* strain. This study. |
| OMF416 | TTCGGCGAGCAGGCGATCCTGC | |
| OMF423 | GTCCATTCGCTGCGCACAGTTGTCG | Verification of Δ*MMAR_1341* strain. This study. |
| OMF424 | TGTTGCCGGTTGCCTGGTATCTGACC | |
| OBJ126 | GGGCCACAACGGGTTATAGC | Verification of Δ*eis* strain. This study. |
| OBJ127 | AACAGTGTCCTGCCGCTGTG | |
| OBJ134 | ATGCGATCGTGGTCGAGGAG | Verification of Δ*MMAR_2836* strain. This study. |
| OBJ135 | CCCGCCAACGAATTGCCTAC | |
| OBJ142 | TTACCGCGTCCGATAGCGTC | Verification of Δ*MMAR_2615* strain. This study. |
| OBJ143 | TAGAGCAGCACGCTGGACAC | |
| OBJ150 | GCACCGAGCAGGAATACTTG | Verification of Δ*eis1* strain. This study. |
| OBJ151 | TTCAGGTTGGTCCAGATGAG | |
| OBJ158 | GGCGATCCTGCAAAGTCCTG | Verification of Δ*mbtK* strain. This study. |

| | | |
|---|---|---|
| OBJ159 | GAAAGACGAGCAGCGGAAGG | |
| OBJ166 | GAATTTGGGTCGGTGACATC | Verification of Δ*MMAR_2744* strain. This study. |
| OBJ167 | CTGCGCTTGTTGGCGTTGTG | |
| OMF100 | GGCTTAAGTATAAGGAGGAAAACATATGACAGACCACGAC CACACCG | Amplifying the *MMAR_1067* gene for pMOP-*1067* plasmid generation. This study. |
| OMF101 | CTCCGTTTAGAGAGGGGTTATACTAGTCAGGCCGGGTTGG CAGC | |
| OMF104 | GGCTTAAGTATAAGGAGGAAAACATATGGATGTCGGTCCG CTGC | Amplifying the *rimJ* gene for pMOP-*rimJ* plasmid generation. This study. |
| OMF105 | CTCCGTTTAGAGAGGGGTTATACTAGTCAGAGCCGGCTGG CTTG | |
| OMF108 | GGCTTAAGTATAAGGAGGAAAACATATGAGCGAAGCGCTG CGCC | Amplifying the *elaA* gene for pMOP-*elaA* plasmid generation. This study. |
| OMF109 | CTCCGTTTAGAGAGGGGTTATACTAGTCATGACTGCGCCG CGGGAC | |
| OMF110 | GGCTTAAGTATAAGGAGGAAAACATATGACTGAAGTTGTCA GCGCCG | Amplifying the *argJ* gene for pMOP-*argJ* plasmid generation. This study. |
| OMF111 | CTCCGTTTAGAGAGGGGTTATACTAGTCATGAGCTGTATG CCGAATTCTCTTC | |
| OMF112 | GGCTTAAGTATAAGGAGGAAAACATATGCATACCGAGGTT CACACGG | Amplifying the *aac* gene for pMOP-*aac* plasmid generation. This study. |
| OMF113 | CTCCGTTTAGAGAGGGGTTATACTAGTTACCAGACGTCTC CCGCG | |
| OMF114 | GGCTTAAGTATAAGGAGGAAAACATATGACCGCCGATCTC GGGC | Amplifying the *rimI* gene for pMOP-*rimI* plasmid generation. This study. |
| OMF115 | CTCCGTTTAGAGAGGGGTTATACTAGTTACCGCGCACCCC CTGGAT | |

| | | |
|---|---|---|
| OMF118 | GGCTTAAGTATAAGGAGGAAAACATATGGCCGAAGTGACCGGCG | Amplifying the *pat* gene for pMOP-*pat* plasmid generation. This study. |
| OMF119 | CTCCGTTTAGAGAGGGGTTATACTAGTCAGCTCACCGCCTCGATCACCT | |
| OMF120 | GGCTTAAGTATAAGGAGGAAAACATATGTCTCGTCACTGGCCGCTGTTCG | Amplifying the *MMAR_4889* gene for pMOP-*4889* plasmid generation. This study. |
| OMF121 | CTCCGTTTAGAGAGGGGTTATACTAGTCAGGCCGCGCGCGAAAC | |
| OMF122 | GGCTTAAGTATAAGGAGGAAAACATATGACGATCTTCGTCATTGATCTGCC | Amplifying the *MMAR_3205* gene for pMOP-*3205* plasmid generation. This study. |
| OMF123 | CTCCGTTTAGAGAGGGGTTATACTAGTCTACAGGGGAAGCGCGCGG | |
| OMF124 | GGCTTAAGTATAAGGAGGAAAACATATGACCCCACAGGCGCGTC | Amplifying the *MMAR_0332* gene for pMOP-*0332* plasmid generation. This study. |
| OMF125 | CTCCGTTTAGAGAGGGGTTATACTAGTTACCGCGGGTCACGCCAC | |
| OMF126 | GGCTTAAGTATAAGGAGGAAAACATATGACCGAGAACATCCGCCGGG | Amplifying the *MMAR_1341* gene for pMOP-*1341* plasmid generation. This study. |
| OMF127 | CTCCGTTTAGAGAGGGGTTATACTAGTCAGCGGGGTCCGGCCAG | |
| OMF132 | GGCTTAAGTATAAGGAGGAAAACATATGACTGCAGTACTGCCCGCA | Amplifying the *eis* gene for pMOP-*eis* plasmid generation. This study. |
| OMF133 | CTCCGTTTAGAGAGGGGTTATACTAGTCTAGAACTCGAAAGCCGTCTCGG | |
| OBJ470 | AGGAGTCCAGCCATATGACTGTGACCCTGTGTAGCC | Amplifying the $eis_{MT}$ gene for pMOP-$eis_{MT}$ plasmid generation. This study. |
| OBJ471 | GCCTGAGCGGTCCCGACTAGTTCAGAACTCGAACGCGGTC | |

| | | |
|---|---|---|
| OMF134 | GGCTTAAGTATAAGGAGGAAAACATATGCACGCCGACGAGGTGGAC | Amplifying the *MMAR_2836* gene for pMOP-*2836* plasmid generation. This study. |
| OMF135 | CTCCGTTTAGAGAGGGGTTATACTAGTCACCACGAGCCAAGCGAGCC | |
| OMF138 | GGCTTAAGTATAAGGAGGAAAACATATGACGGAGATGCCCGACGGTAT | Amplifying the *MMAR_2615* gene for pMOP-2615 plasmid generation. This study. |
| OMF139 | CTCCGTTTAGAGAGGGGTTATACTAGTCAGCTCTGGGATCTCGAGCGGT | |
| OMF142 | GGCTTAAGTATAAGGAGGAAAACATATGAGCACCAACATCCGCGTTC | Amplifying the *eis1* gene for pMOP-*eis1* plasmid generation. This study. |
| OMF143 | CTCCGTTTAGAGAGGGGTTATACTAGTCTAGAAGAAGAACCCGGCGTGGG | |
| OMF140 | GGCTTAAGTATAAGGAGGAAAACATATGGCGGGCCATCCGGGC | Amplifying the *mbtK* gene for pMOP-*mbtK* plasmid generation. This study. |
| OMF141 | CTCCGTTTAGAGAGGGGTTATACTAGTCATCGGCGTTGCCGGTCACG | |
| OMF146 | GGCTTAAGTATAAGGAGGAAAACATATGACTGCCGACAGGACCGG | Amplifying the *MMAR_2744* gene for pMOP-*2744* plasmid generation. This study. |
| OMF147 | CTCCGTTTAGAGAGGGGTTATACTAGTCAAACGGGGTCGGTGATCTCGTC | |
| OBJ529 | AGGAGTCCAGCCATATGTTCCCCGGAGCCGTAATC | Amplifying the *papA5* gene for pMOP-*papA5* plasmid generation. This study. |
| OBJ530 | GCCTGAGCGGTCCCGACTAGTTCATTCCATGAACAACCCGTACTCTG | |
| OBJ426 | AGGAGTCCAGCCATATGAGTGAAGCCGACACGC | Amplifying the *mbtK1* gene for pMOP-*mbtK1* plasmid generation. This study. |
| OBJ427 | GCCTGAGCGGTCCCGACTAGTTCAGCTGCGTAGTCCGGG | |
| OBJ476 | GAGGGCGGCATCGCCGGCCGATTCGGC | Site-directed mutagenesis primers *eisY128A*. This study. |

| | | |
|---|---|---|
| OBJ477 | GCCGAATCGGCCGGCGATGCCGCCCTC | |
| OBJ482 | CGTGTCATCGCCGCCCCCGACGTCAAC | Site-directed mutagenesis primers *mbtKE161A.* This study. |
| OBJ483 | GTTGACGTCGGGGGCGGCGATGACACG | |
| pMOP-Seq-F | GCCTTTGAGTGAGCTGATAC | Forward sequencing primer for pMOP expression plasmid. |
| OMF101 | CTCCGTTTAGAGAGGGGTTATACTAGTCAGGCCGGGTTGGCAGC | Reverse primer for verification of *pMMAR_1067* plasmid. This study. |
| OMF103 | CTCCGTTTAGAGAGGGGTTATACTAGTTAAAGCACCAACAGCATCCGG | Reverse primer for verification of *pargA* plasmid. This study. |
| OMF105 | CTCCGTTTAGAGAGGGGTTATACTAGTCAGAGCCGGCTGGCTTG | Reverse primer for verification of *primJ* plasmid. This study. |
| OMF107 | CTCCGTTTAGAGAGGGGTTATACTAGTCAGTCGAGCAGCACCGTGG | Reverse primer for verification of *pemp1* plasmid. This study. |
| OMF109 | CTCCGTTTAGAGAGGGGTTATACTAGTCATGACTGCGCCGCGGGAC | Reverse primer for verification of *pelaA* plasmid. This study. |
| OBJ58 | GCCTGAGCGGTCCCGACTAGTAATCGTCATGAGCTGTATGC | Reverse primer for verification of *pargJ* plasmid. This study. |
| OMF113 | CTCCGTTTAGAGAGGGGTTATACTAGTTACCAGACGTCTCCCGCG | Reverse primer for verification of *paac* plasmid. This study. |
| OMF115 | CTCCGTTTAGAGAGGGGTTATACTAGTTACCGCGCACCCCCTGGAT | Reverse primer for verification of *primI* plasmid. This study. |
| OMF119 | CTCCGTTTAGAGAGGGGTTATACTAGTCAGCTCACCGCCTCGATCACCT | Reverse primer for verification of *ppat* plasmid. This study. |
| OMF121 | CTCCGTTTAGAGAGGGGTTATACTAGTCAGGCCGCGCGCGAAAC | Reverse primer for verification of *pMMAR_4889* plasmid. This study. |
| OMF123 | CTCCGTTTAGAGAGGGGTTATACTAGTCTACAGGGGAAGCGCGCGG | Reverse primer for verification of *pMMAR_3205* plasmid. This study. |

| | | |
|---|---|---|
| OMF125 | CTCCGTTTAGAGAGGGGTTATACTAGTTACCGCGGGTCAC GCCAC | Reverse primer for verification of *pMMAR_0332* plasmid. This study. |
| OMF127 | CTCCGTTTAGAGAGGGGTTATACTAGTCAGCGGGGTCCGG CCAG | Reverse primer for verification of *pMMAR_1341* plasmid. This study. |
| OMF133 | CTCCGTTTAGAGAGGGGTTATACTAGTCTAGAACTCGAAA GCCGTCTCGG | Reverse primer for verification of *peis* plasmid. This study. |
| OBJ372 | GCCTGAGCGGTCCCGACTAGTTCACCACGAGCCAAGCGA GCC | Reverse primer for verification of *pMMAR_2836* plasmid. This study. |
| OBJ364 | AATGCTCATCCCGACGATCT | Reverse primer for verification of *pMMAR_2615* plasmid. This study. |
| OBJ366 | TCAGATCGAGTCCGAACAGG | Reverse primer for verification of *peis1* plasmid. This study. |
| OMF141 | CTCCGTTTAGAGAGGGGTTATACTAGTCATCGGCGTTGCC GGTCACG | Reverse primer for verification of *pmbtK* plasmid. This study. |
| OBJ370 | ACATGCTCGGGATCGACATA | Reverse primer for verification of *pMMAR_2744* plasmid. This study. |
| OMF049 | GTTGGACTCAAGACGATAGTTACCGGATAAG | Δ*mbtK /* p*rimI* cross-complement verification. This study. |
| OBJ62 | GCCTGAGCGGTCCCGACTAGTCATTTACCGCGCACC | |
| OMF049 | GTTGGACTCAAGACGATAGTTACCGGATAAG | Δ*mbtK /* p*pat* cross-complement verification. This study. |
| OBJ64 | GCCTGAGCGGTCCCGACTAGTCGATCAGCTCACCGCCTC | |
| OMF049 | GTTGGACTCAAGACGATAGTTACCGGATAAG | Δ*mbtk /* p*MMAR_3205* cross-complement verification. This study. |
| OBJ68 | GCCTGAGCGGTCCCGACTAGTGGCCTACAGGGGAAG | |
| OMF049 | GTTGGACTCAAGACGATAGTTACCGGATAAG | Δ*mbtK /* p*eis* cross-complement verification. This study. |
| OMF133 | CTCCGTTTAGAGAGGGGTTATACTAGTCTAGAACTCGAAA GCCGTCTCGG | |

| | | |
|---|---|---|
| OBJ471 | GCCTGAGCGGTCCCGACTAGTCAGAACTCGAACGCGGTC | Δ*mbtK* / p*eis*$_{MT}$ cross-complement verification. This study. |
| OBJ366 | TCAGATCGAGTCCGAACAGG | Δ*mbtK* / p*eis1* cross-complement verification. This study. |
| OBJ530 | GCCTGAGCGGTCCCGACTAGTCATTCCATGAACAACCCGTACTCTG | Δ*mbtK* / p*papa5* cross-complement verification. This study. |
| Kan For | CGAGGCCGCGATTAAATTC | Confirmation of kanamycin resistance gene presence. This study. |
| Kan Rev | AAACTCACCGAGGCAGTTC | |

**Dataset S1**

Proteomics data supporting this manuscript, including A. unprocessed data, Processed B. Cell associated protein levels and C. Secreted protein levels. D. levels of N-terminal acetylation.

**Supplementary Methods:**

**<u>Growth & Maintenance of Bacterial Strains:</u>** *Mycobacterium marinum* strains were derived from the *Mycobacterium marinum* M strain (ATCC BAA-535). *M. marinum* strains were maintained in Middlebrook 7H9 (Sigma-Aldrich) defined broth supplemented with 0.5% glycerol and 0.1% Tween-80. For solid media, *M. marinum* strains were maintained on Middlebrook 7H11 agar (Sigma-Aldrich) supplemented with 0.5% glucose and 0.5% glycerol. 7H9 and 7H11 media were additionally supplemented with 20 µg/mL kanamycin (IBI Scientific), 50 µg/mL hygromycin, or 60 µg/mL X-gal, where appropriate. Where noted and appropriate, some cultures were supplemented with ferric mycobactin J (Allied Monitor). Strains were grown at 30°C. For *Escherichia coli* strains, *E. coli* DH5α and supplemented with 50 µg/mL kanamycin, or 200 µg/mL hygromycin, where appropriate. *E. coli* were grown at 37°C. All strains in this study are listed in Table S1.

**<u>Generation of *M. marinum* Strains:</u>** Unmarked *Mycobacterium marinum* deletion strains created were derived from the *Mycobacterium marinum* M strain. Genetic deletions utilized within this study were created using allelic exchange, as previously described (5, 6, 10, 11). ~ 1,500 bp upstream and downstream of a given gene's annotated open reading frame were amplified via PCR. Amplified regions were cloned into the p2NIL vector [(Addgene plasmid number 20188; a gift from Tanya Parish, (7)] by FastCloning (12) as previously described (11, 13). The p2NIL vector, now containing regions flanking the gene of interest, was digested with PacI restriction enzyme (New England Biolabs) and treated with Antarctic Phosphatase (New England Biolabs). pGOAL19 vector [Addgene plasmid number 20190; a gift from Tanya Parish (7)] was also digested with PacI restriction enzyme, and heat inactivated at 65°C. PacI digested pGOAL19 was ligated into the PacI, Antarctic Phosphatase treated p2NIL vector as previously described (11, 13). The resulting plasmid was quantified on a NanoDrop Microvolume Spectrophotometer (Thermo Fisher). 2 µg of plasmid was irradiated with 0.1 J/cm2 UV light in a CL-1000 UV crosslinker (UVP), followed by electroporation into 500 µl of electrocompetent *M. marinum* cells using a Gene Pulser Xcell Electroporation System (Bio-Rad). *M. marinum* electrocompetent cells were prepared as described (11, 13). After electroporation, competent cells were recovered in 2 mL of 7H9 defined broth supplemented with 0.1% Tween-80 and incubated at 30°C overnight. Cells were collected by centrifugation and plated on 7H11 agar (Sigma) plates supplemented with 20 µg/ml kanamycin, 60 µg/ml X-Gal, and oleic acid-albumin-dextrose-catalase (OADC). Merodiploid colonies stemming from the aforementioned 7H11 agar plates were selected and grown in 5 mL of 7H9 defined broth supplemented with 0.1% Tween-80. 300 µl of culture was collected and lysed using zirconia disruption beads (RPI) and a BioSpec Mini-Beadbeater-24 followed by centrifugation. 1 µl of lysed bacteria was utilized in PCR experiments to confirm deletion or complementation of each strain. PCR productions were visualized in TAE agarose gels stained with ethidium bromide (VWR) on a Gel Doc EZ Imager (Bio-Rad), with the size of the PCR product being indicative of wild-type or knockout of a given gene. Genetic deletions were confirmed via targeted Sanger DNA sequencing at the Notre Dame Genomics and Bioinformatics Facility, or at Plasmidsaurus.

**Generation of Complementation Plasmids:** Complementation plasmids were generated by amplifying a gene of interest from *M. marinum* M strain genomic DNA using the primers listed in Table S3. Some genes were cloned directly into the pMOP vector via FastCloning (12). Some genes were first cloned into the pUC19 vector, and the gene of interest was subsequently moved to the pMOP vector via restriction cloning. Briefly, a gene of interest was isolated from the pUC19 plasmid via NdeI (New England Biolabs) and SpeI (New England Biolabs) restriction digestion, followed by gel purification. The purified NdeI and SpeI digested gene of product of interest was then ligated into NdeI and SpeI digested pMOP vector to assemble the completed pMOP-*gene* plasmid. Complementation plasmids created were confirmed via targeted Sanger DNA sequencing at the Notre Dame Genomics and Bioinformatics Facility, or at Plasmidsaurus.

**Laboratory Growth Kinetics of *M. marinum* Strains:** Mycobacterial growth kinetics of strains indicated in Figure S4A were performed as follows: Mycobacterial strains indicated were grown in a 5 mL volume of 7H9 defined broth supplemented with 0.1% Tween-80 for approximately 3 days. Bacteria present in the culture were approximated by optical density at $OD_{600}$. Mycobacteria were sub-cultured into a 25 mL volume of 7H9 defined broth supplemented with 0.1% Tween-80 at an $OD_{600}$ of 0.2. 25 mL cultures were grown at 30°C, shaking, for 7 days. Bacterial growth was assessed every 24 hours over the course of 7 days by $OD_{600}$ reading. Where indicated, some cultures in Figure S4A were supplemented with 2 µg/mL ferric mycobactin J (Allied Monitor).

**Hemolysis Assays:** Hemolysis assays were performed exactly as described in (11).

**Cytotoxicity Macrophage Infections:** RAW 264.7 cells (ATCC, TIB-71) were cultured and passaged as described (14). RAW 264.7 cells were seeded in 200 µL DMEM supplemented by 10% FBS per well at $2.5x10^5$ cells/mL in a clear 96-well plate (Thermo Fisher). After 24 hours of growth, RAW 264.7 cells were infected with bacteria at an estimated MOI of 5 ($2.5 \times 10^5$ cells/mL) in technical triplicate. 2 hours post-infection (hpi), extracellular bacteria were killed by gentamycin addition (RPI Corporation) at 100 µg/mL for 2 hours. Then, gentamycin was removed by washing three times with sterile 1x PBS and adding fresh DMEM plus 10% FBS. 24 hpi, medium was changed by 50 µL solution of 1x PBS containing EthD-1 (1 µL/mL) (Live/Dead viability/cytotoxicity kit; Life Technologies) and Hoechst 33342 (0.33 µg/mL) (Thermo Fisher). Cells were incubated for an additional 30 min at 37°C. Cells were imaged as described previously (14), except that the DAPI filter was used instead of the green fluorescent protein filter. Five images were taken per well, and ImageJ software was then used to quantify the number of red cells per image, except that a minimum of 4 pixels was set to count the number of particles. A minimum of three independent biological replicates, each containing three technical replicates, was used for statistical analysis.

***Galleria mellonella* Infections:** *Galleria mellonella* (Greater Wax Moth larvae, “wax worms”) were purchased (waxworms.net) and sorted to select for live, non-melanized worms with a mass of greater than 200 mg. Following growth in 7H9 media for approximately three days at 30°C, *M. marinum* cells were collected by centrifugation, washed three times with 1x PBS, and were resuspended in a final volume of 1mL. *M. marinum* were syringed with a 27G ½” needle (Exel International, Salaberry-de-Valleyfield, Quebec, Canada) and allowed to settle for 30 minutes. The optical density ($OD_{600}$) of the single cell suspension was enumerated, the *M. marinum* were collected by centrifugation and resuspended in 200 µL to generate a concentration of $1x10^7$ / 5 µL. 5 µL of the single cell suspension was injected into each *Galleria mellonella* larvae as performed in (2, 15).

*Galleria mellonella* larvae were swabbed with 50% ethanol prior to injection of 5μL of *M. marinum* in the worm's back right-most proleg using a 10μL syringe (Hamilton Company, Reno, NV). Worms were injected in triplicate in groups of 10, resulting in 30 worms used per strain per biological replicate. After injection, worms were incubated at 30°C for the infection over 7 days. Any worms irresponsive to physical touch and melanized were declared deceased. Any larvae proceeding to their pupal stage during the infection time course were removed and disposed of appropriately to avoid metamorphosis into the Greater Wax Moth. Statistical analysis was performed as in (2).

**Thin-Layer Chromatography:** Thin-layer chromatography experiments to observe mycobacterial lipids were performed exactly as described in (6).

**Site-Directed Mutagenesis:** Site-directed mutagenesis experiments were performed according to the QuikChange II Site-Directed Mutagenesis Kit user manual (Agilent) exactly as described previously (16).

**Expression and purification of 6×His tagged proteins:** The *eis, eisY128A*, *mbtK*, *mbtKE161A* genes were amplified from the pMOPS plasmid. The $eis_{MT}$ (*ERDMAN_4236*) gene was amplified from the *M. tuberculosis* Erdman genomic DNA using Phusion polymerase (NEB) and primers (IDT) listed in Table S2. Cloning vectors pET15b (for *eis, eisY128A* and *Rv2416c*) and pET28b-MBP-TEV (Addgene plasmid # 69929 for *mbtK* and *mbtKE161A*) were prepared for fast cloning by PCR amplification using Phusion polymerase, HF buffer and the primers (Table S2) (17). The vector and inserts were mixed, treated with Dpn1(12), and then introduced into chemically competent DH5α *E. coli* cells. These cells were plated on LB agar plates containing 200 μg/mL ampicillin (for pET15b) or 50 μg/mL Kanamycin (for pET28b-MBP-TEV) and incubated at 37°C. Individual colonies were purified, grown overnight in LB broth in presence of antibiotic at 37°C, followed by plasmid extraction (Bioneer). Plasmids were evaluated by restriction digestion with NdeI/SpeI enzymes (for pET15b constructs) and NdeI/AflII (for pET28b-MBP-TEV constructs). *rimI* from *Salmonella typhimurium* was cloned for *in vitro* expression by amplification from the construct pET23a-RimIST (17) using primers OMF088, OMF089. *rimI* from *S. typhimurium* (*rimIST*) was amplified for FastCloning (12) using the primers OMF310, OMF311. pET28MBP (18) was amplified for FastCloning using Phusion polymerase, HF buffer and the primers OMF033, OMF034. Plasmids were evaluated by restriction digestion with NdeI/SpeI enzymes. All resulted plasmids were confirmed by targeted DNA sequencing by the Genomics and Bioinformatics Core at the University of Notre Dame or by Plasmidsaurus.

The expression constructs were introduced into BL21 Star *E. coli* cells. Colonies were used to start a 5 mL starter culture, which was used to inoculate 200 mL of auto-induction media (1:100 ratio) (18) in the presence of 200 µg/mL ampicillin (pET15b constructs) or 50 µg/mL kanamycin (pET28b-MBP-TEV constructs). Cultures were incubated in shaking condition at 30°C for 24h (for *eis, eisY128A, and* $eis_{MT}$) and 20h (for *mbtK, mbtKE161A*, and $rimI_{ST}$) followed by the bacterial cell harvest as mentioned in our previous report (13). Bacterial cells were lysed and proteins were eluted by following the aforementioned methodology (13). Final eluted proteins were quantified using MICRO BCA protein assay kit (Thermo scientific) with 6×His-$Eis_{MM}$ at 6.8 µg/µL, 6×His-Eis $Y128A_{MM}$ at 6.94 µg/µL, 6×His-$Eis_{MT}$ at 3.02 µg/µL, 6×His-$MbtK_{MM}$ at 13.4 µg/µL, 6×His-MbtK $E161A_{MM}$ at 11.5 µg/µL, 6×His-$rimI_{ST}$ at 7.5 µg/µL.

**Proteolytic Tag Removal:** Proteins expressed in the pET28MBP vector feature a Tobacco Etch Virus (TEV) protease site (ENLYFQ/G) downstream of the affinity tags allowing for their proteolytic removal. Using an 6×His affinity tagged protease allows for protease, cleaved tag and uncleaved protein to be removed by subtractive affinity chromatography over Ni-NTA resin. 6×His-tagged Tobacco Etch Virus (TEV) protease was expressed from the plasmid pRK793 (Addgene plasmid # 8827 (19) a gift of David Waugh) by autoinduction and purified as above.

Elution fractions 6×His-RimI were pooled, mixed with 50 µL of purified TEV and loaded into 3,000 or 7,000 MWCO dialysis tubing. Mixture was dialyzed at 4°C overnight against cleavage buffer [20 mM HEPES pH 7.0, 10% glycerol, 300 mM NaCl, 5 mM β-mercaptoethanol]. Dialysis tubing was transferred to reductant-free buffer [20 mM HEPES pH 7.0, 10% glycerol, 300 mM NaCl] and dialyzed at 4°C for 5 hours. Cleaved proteins were removed from the dialysis tube, mixed with buffer-equilibrated Ni-NTA resin and batch incubated at 4°C for 1 hour. Protein resin slurries were loaded into empty PD-10 columns and the flow through was collected. Resin was washed with 2 column volume of buffer which was added to the flow through. Pooled flow-through plus wash was concentrated by ultra-filtration in a 3000 MWCO Amicon (Millipore) and quantified by BCA assay (Pierce). Concentrated proteins were aliquoted into PCR tubes, flash-frozen in liquid nitrogen and stored at -80°C.

**5,5-dithio-bis-(2-nitrobenzoic acid) (DTNB) Assays:** The 5,5′-dithiobis-(2-nitrobenzoic acid) (DTNB) was performed as previously described with some modifications (20). Briefly, purified protein (~7-10 µg) was combined with synthetic peptides (300 µM) and Ac-CoA (300 µM) in acetylation buffer (50 mM Tris-HCl pH 8.5, 200 mM NaCl, and 2 mM EDTA) at 37 °C. After 30 minutes, the reactions were halted with quenching buffer (3.2 M guanidinium-HCl, 100 mM sodium phosphate dibasic pH 6.8). N-terminal acetylation process transfers the acetyl group from Ac-CoA to the N-termini of peptides, which exposes a thiol-group on CoA. The CoA production was assessed by introducing DTNB (2 mM final, dissolved in 100 mM sodium phosphate dibasic pH 6.8 and 10 mM EDTA) to the quenched reactions. DTNB readily react with the thiol that yields 2-nitro-5-thiobenzoate ($TNB^{-}$) that will ionize to $TNB^{2-}$, which is easily quantified by measuring the absorbance at 412 nm. Background absorbance was determined in negative controls (enzyme added after quenching buffer) and subtracted from the absorbance in each reaction. $TNB^{-}$ production was quantified assuming $\varepsilon = 13.7 \times 10^3\ M^{-1}\ cm^{-1}$.

**Proteomics: (Sample prep, Instrument, PEAKS search):**

LC-MS pure reagents (water, ethanol, acetonitrile, and methanol) were purchased from J.T. Baker (Radnor, PA). Iodoacetamide (IAA) was purchased from MP Biomedicals (Solon. OH). All other reagents are from Sigma-Aldrich (St. Louis, MO) unless specified. S-Trap mini devices were from Protifi (Huntington, NY). Trypsin Gold was purchased from Promega (Madison, WI). Hydrophilic–lipophilic balance (HLB) solid phase extraction (SPE) cartridges (1 cc/10 mg) from Waters (Milford, MA) were used to desalt peptide samples prior to analysis on an Evosep One (Odense, Denmark) and timsTOF Pro from Bruker Scientific (Billerica, MA). Protein and peptide identification and label-free quantitation was performed using the PEAKS Online X search engine from Bioinformatics Solutions Inc (Waterloo, ON) (build 1.4.2020-10-21_171258). MALDI-MS was performed on an Ultraflextreme from Bruker Scientific (Billerica, MA).

***MALDI*:** For each sample, 1μL of sample was spotted on an MTP 384 ground steel target, left to dry, and followed by 1μL of saturated α-cyano-4-hydroxycinnamic acid in a solution of 0.1% formic acid in 50% acetonitrile. Samples were analyzed on a Bruker Ultraflextreme MALDI mass spectrometer. 8,000 shots were summed at 500Hz laser frequency with a mass range of 400-4,000 m/z. For detector gain, reflector was set to 32x (2729V).

***LC-MS*:** N-Acetoxy-D3-succinamide was synthesized as described by Thompson et al [1]. Briefly, 5.0g of $D_6$ acetic anhydride was incubated with 1.77g of N-hydroxysuccinamide in a round-bottom flask at room temperature overnight, with aluminum foil loosely wrapped over the mouth. The following day, aluminum foil was removed and stirring continued to begin evaporation of excess acetic acid/anhydride. Contents were dried using rotary evaporation without heat (~1mbar vacuum), and rinsed with 50mL of hexanes or enough to submerge product. Three more wash and dry steps were performed. Exactly 7.0mL of anhydrous acetonitrile was added to fully dissolve the dried product. From this solution, 141.9μL aliquots were made into 1.5mL microcentrifuge tubes, producing 50mg aliquots of $d_3$-acetoxy-NHS ester product.

Cell-associated and secreted mycobacterial protein fractions were generated from *M. marinum* as described in Cronin et al., 2022 (10). Cell-associated and secreted cell lysate samples were prepared for LC-MS analysis as described in Saelens et al and Collars et al (6, 21). 50µg of each sample was precipitated in at least 7x volume of cold acetone for 2 hours, followed by centrifugation (10 minutes at 12,000xg), decanting, and drying. Samples were resuspended in 100mM triethylammonium bicarbonate (TEAB), 5% sodium dodecyl sulfate (SDS), and 100mM tris(2-carboxyethyl)phosphine (TCEP). Samples were heated for 10 minutes at 95˚C, then added to 100mM IAA for 30 minutes in dark. Samples were acidified with phosphoric acid to a final concentration of 1.2% and volume of 50µL, then flocculated with 350µL of a solution containing 90% methanol and 10% 1M TEAB (Binding Buffer). Samples were passed through S-Trap Mini filters and followed by 80µL of Binding Buffer and 80µL of 1:1 methanol/chloroform solution.

Previous 50mg NHS ester aliquot was resuspended in 518µL of 10% methanol in acetonitrile. 75µL of NHS esters were added to each sample filter, followed by 30 minutes of incubation at 25˚C. Samples were spun down and a second addition and incubation of 75µL esters was performed. Samples were then quenched with two washes of 150µL 50mM ammonium bicarbonate in 90% methanol. A new collection tube was replaced for each sample, and 1µg of trypsin in 80µL of 100mM TEAB buffer was added. Samples were wrapped in Parafilm to prevent evaporative loss and incubated at 37˚C for 12 hours. Digested peptides were spun through the filter, followed by two 80µL elutions with 0.1% formic acid in water, and one 80µL elution with 0.1% formic acid in 50% acetonitrile. Eluted peptides were vacuum concentrated for 20 minutes (to remove acetonitrile), and then desalted using 1cc/10mg HLB SPE cartridges following manufacturer's specifications.

Desalted peptides were vacuum concentrated to dryness and resuspended in 0.1% formic acid and water, to 1 mg/mL concentration. 500ng of each sample was injected in triplicate onto a Bruker nanoElute 2 and timsTOF Pro LC-MS system. Each sample was prepared in biological triplicate and technical triplicate. 120min gradient LC methods on a 150 µm x 250 mm PepSep

XTREME column with C18 ReproSil AQ stationary phase at 1.5µm particle size, 100Å pore size. Nano-ESI was used as the method of ionization, with a spray voltage of 1700V. MS was set to Parallel Accumulation, Serial Fragmentation Data Dependent Mode (PASEF-DDA) with a mass range of 100–1700 m/z, ion mobility range of 0.6–1.6 v*s/cm2, and ramp and accumulation times of 100 ms. Each precursor consisted of 10 PASEF ramps for a cycle time of 1.17 s. Precursors were filtered to contain only charges from 2 to 5. MS/MS collision energy settings were set to ramp from 20 eV at 0.6 ion mobility to 70 eV at 1.6 ion mobility. Instrument tune parameters were set to default for proteomic studies with the following differences: quadrupole low mass set to 20 m/z, focus pre-TOF pre-pulse storage set to 5 ms.

Protein and peptide identification and label-free quantitation were performed using the PEAKS Online X search engine. Search database used was the *M. marinum* proteome from Mycobrowser (v4 release) with the following adjustments; the initiator codon for a significant number of mycobacterial protein entires is not iMet. Mycobrowser FASTAs incorrectly translates these as coded (e.g. GTG->Val). These were recoded manually to Met. Search settings were set to manufacturer defaults unless specified. 3 missed cleavages were allowed, at a semi-specific search. Fixed modification included carbamidomethylation of cysteine, while variable modifications included protein $H_3$ and $D_3$ N-terminal acetylation, $D_3$ acetylation of Lys,deamidation of asparagine and glutamine, pyroglutamic acid from glutamine and glutamic acid, and oxidation of methionine. All peptide-spectrum matches were filtered to a 1% false discovery rate. Cell-associated samples were normalized by total ion current (TIC). Raw data is available through MassIVE and The Proteome Exchange at ftp://MSV000095200@massive.ucsd.edu with Identifier MSV000095200 PXD053528.

### Proteomics Data Analysis

Protein Level Analysis: Protein database search files were exported from PEAKS (SI Data) and contaminants were filtered out. Data were processed using R (v4.3.2), and the code can be found on github (https://github.com/Champion-Lab/3692). The pellet and supernatant fractions were analyzed independently. The workflow was adapted from Hutchings *et al*. (22). Briefly, protein areas were loaded into Qfeatures (v1.10.0)(23). log transformed and median normalized. Technical replicates (n=3) were averaged to obtain a quantitative value for each biological replicate (n=3). Differential expression analysis was performed with these values using limma (v3.56.2) and the ebayes() function (24) to calculate fold change and significance, along with a Benjamini-Hochburg (B-H) correction (25) comparing WT to Δ*mbtK,* Δ*mbtK/*p*mbtK*, *and* Δ*mbtK/*p*eis* individually. Missing values were excluded from these comparisons, but added to the downstream plots as either infinite or negative infinite fold changes. Log fold-change was plotted against the WT expression levels (log transformed and normalized, Fig. S5). Proteins that had a significant fold change between WT and Δ*mbtK* were highlighted in all plots, along with other annotated Mbt proteins as large points with labels, colored by whether their B-H adjusted *p-value* was less than 0.05. Error bars represent 95% confidence intervals calculated from the B-H corrected *p-values (26)*. Supernatant plots also include labeled proteins that were significantly changing in the pellet.

**Peptide Level Acetylation Analysis:**

Peptide database search files were exported from PEAKS (SI Data) and contaminants were filtered out. Data were processed using R (v4.3.2), and the code can be found on github (https://github.com/Champion-Lab/3692). Briefly, peptides with N-terminal acetylation modifications (light: +42.01, heavy: +45.03) and that start at amino acid position 1 or 2 in the protein sequence were analyzed. The 'best flier' light acetylated N-terminal peptide for each protein was identified by taking the peptidoform with the highest mean area across all injections. This best flier and its heavy cognate peptide were compared to generate a percent acetylation (%acet) value for each protein for each injection [(light / light + heavy) * 100]. These values were averaged for each strain (WT, Δ*mbtK,* Δ*mbtK/*p*mbtK*, *and* Δ*mbtK/*p*eis*). Proteins where %acet of the Δ*mbtK* was ≤ WT and Δ*mbtK/*p*mbtK* were identified as potential N-terminal acetylation targets of *mbtK* (Dataset S1) and %acet was plotted for each strain for these proteins (Fig. S11). Additional measurements of precision are available in the SI.
